# Neurogenetic and genomic approaches reveal roles for Dpr/DIP cell adhesion molecules in *Drosophila* reproductive behavior

**DOI:** 10.1101/2020.10.02.323477

**Authors:** Savannah G Brovero, Julia C Fortier, Hongru Hu, Pamela C Lovejoy, Nicole R Newell, Colleen M Palmateer, Ruei-Ying Tzeng, Pei-Tseng Lee, Kai Zinn, Michelle N Arbeitman

**Author notes:** co-first authors (Savannah G Brovero, Julia C Fortier, Hongru Hu, Pamela C Lovejoy, Nicole R Newell, Colleen M Palmateer, Ruei-Ying Tzeng).

## Abstract

*Drosophila* reproductive behaviors are directed by *fruitless* neurons (*fru P1* isoforms). A reanalysis of genomic studies shows that genes encoding *dpr* and *DIP* Immunoglobulin superfamily (IgSF) members are expressed in *fru P1* neurons. Each *fru P1*and *dpr/DIP* (*fru P1* ∩ *dpr/DIP*) overlapping expression pattern is similar in both sexes, with dimorphism in neuronal morphology and cell number. Behavioral studies of *fru P1* ∩ *dpr/DIP* perturbation genotypes point to the mushroom body functioning together with the lateral protocerebral complex. Functionally, we find that perturbations of sex hierarchy genes and *DIP-ε* changes sex-specific morphology of *fru P1* ∩ *DIP-α* neurons. A single-cell RNA-seq analysis shows that the *DIPs* have high expression in a restricted set of *fru P1* neurons, whereas the *dprs* are expressed in larger set of neurons at intermediate levels, with a myriad of combinations.

## Introduction

A current goal of neuroscience research is to gain molecular, physiological and circuit-level understanding of complex behavior. *Drosophila melanogaster* reproductive behaviors are a powerful and tractable model, given our knowledge of the molecular-genetic and neural anatomical basis of these behaviors in both sexes. Small subsets of neurons have been identified as critical for all aspects of reproductive behaviors—these neurons express *Drosophila* sex hierarchy transcription factors encoded by *doublesex* (*dsx*) and *fruitless* (*fru; fru P1* transcripts spliced by sex hierarchy; **Figure 1A**) (reviewed in Dauwalder 2011; Yamamoto *et al*. 2014; Andrew *et al*. 2019; Leitner and Ben-Shahar 2020). It is clear that these *dsx*- and *fru P1*-expressing neurons are present in males and females in similar positions, and arise through a shared developmental trajectory (Ren *et al*. 2016), even though these neurons direct very different behaviors in males and females. Males display an elaborate courtship ritual that includes chasing the female, tapping her with his leg, and production of song with wing vibration (reviewed in Greenspan and Ferveur 2000). The female decides whether she will mate and then, if mated, she displays post-mating behaviors that includes egg laying, changes in diet, and changes in her receptivity to courtship (see Laturney and Billeter 2014; Aranha and Vasconcelos 2018; Newell *et al*. 2020).

**Figure 1.**
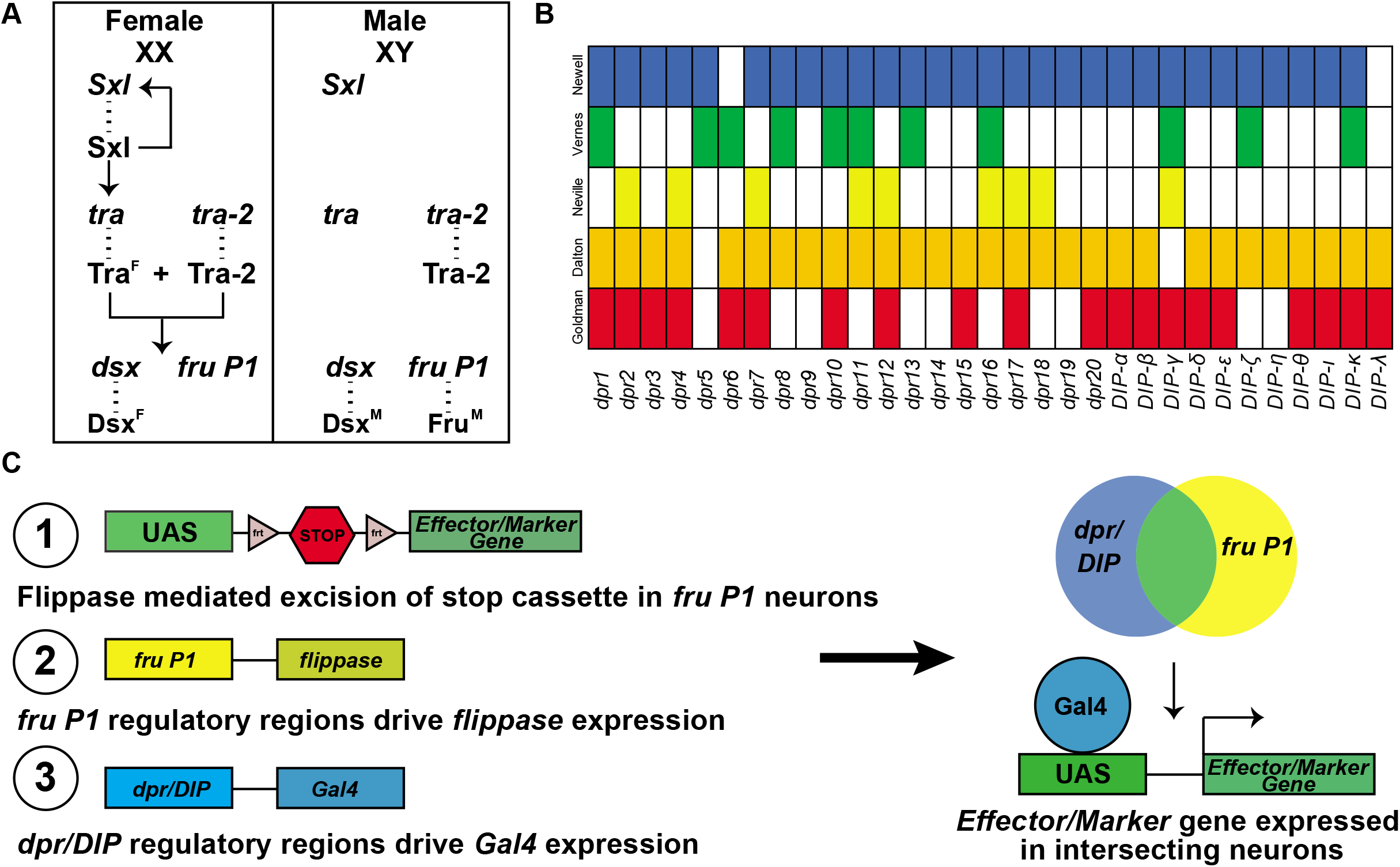
Overview of sex hierarchy and experimental design. **A**) The Drosophila somatic sex determination hierarchy is an alternative pre-mRNA splicing cascade. The presence of two X chromosomes in females results in splicing of *Sxl* pre-mRNA, such that functional Sxl is produced. Sxl regulates *Sxl* and *tra* pre-mRNA splicing, resulting in continued production of functional Sxl and Tra in females. Tra and Tra-2 regulate the pre-mRNA splicing of *dsx* and *fru P1* in females, whereas in males *dsx* and *fru P1* are spliced by the default pre-mRNA splicing pathway. The sex-specific splicing results in production of sex-specific Dsx and Fru transcription factors. *dsx* regulates sex differences that lead to both dimorphic behavior and gross anatomical morphological differences, whereas *fru P1* regulates sex differences that lead to dimorphic behaviors. **B**) Previous genome-wide studies found that *dpr/DIPs* are regulated downstream of *fru P1*, Fru^M^, and/or are expressed in *fru P1*-expressing neurons (Goldman and Arbeitman 2007; Dalton *et al*. 2013; Neville *et al*. 2014; Vernes 2014; Newell *et al*. 2016). **C**) A genetic intersectional strategy was used to express marker or effector genes in *fru P1* ∩ *dpr/DIP* neurons. This strategy takes advantage of the two-component Gal4/UAS expression system, and flippase-mediated removal of a stop cassette within an expression vector. Expression of the marker/effector gene requires both removal of the stop cassette via *fru P1-flippase* (*flp*) expression and expression of Gal4 via *dpr/DIP* regulation. Therefore, only neurons that express both *fru P1* and one of the *dpr/DIPs* have expression of the effector or marker (shown on right).

Sex differences in the nervous system that contribute to reproductive behaviors include dimorphism in *dsx* and *fru P1* neuron number, connectivity, and physiology, with the molecules and mechanisms that direct these differences beginning to be elucidated. Here, through a systematic reanalysis of several genomic studies we show that a set of cell adhesion molecules that are members of the immunoglobulin superfamily (IgSF) are regulated by male-specific Fru (Fru^M^) or are expressed in *fru P1* neurons (**Figure 1B**) (Goldman and Arbeitman 2007; Dalton *et al*. 2013; Neville *et al*. 2014; Vernes 2014; Newell *et al*. 2016). This led us to investigate the role of the Dpr (*<u>d</u>efective <u>p</u>roboscis extension <u>r</u>esponse*) and DIP (*<u>D</u>pr <u>i</u>nteracting <u>p</u>rotein*) IgSF cell adhesion molecules in *fru P1* neurons and the functions of the neurons in which they are expressed for courtship behavior. Sex-specific splicing of transcripts produced from the *fru P1* promoter results in production of Fru^M^ transcription factors that are members of the BTB-zinc finger family, but no female-specific transcription factors (**Figure 1A**) (Ito *et al*. 1996; Ryner *et al*. 1996). The other *fru* transcripts are not sex-specifically spliced and provide essential functions (Anand *et al*. 2001). In addition to the genomic studies, our work showed that *dpr1*, the founding member of the *dpr* family (Nakamura *et al*. 2002) has a role in gating the timing of the steps that comprise the male courtship ritual (Goldman and Arbeitman 2007). The Dpr and DIP proteins are classified as cell-adhesion molecules, given that they are transmembrane proteins that contain extracellular Ig domains, with short cytoplasmic tails. The Dpr proteins have two extracellular Ig domains, whereas DIPs have three Ig domains (reviewed in Zinn and Ozkan 2017; Sanes and Zipursky 2020). The finding that cell adhesion molecules are regulated by Fru^M^ fit well with studies that showed that there are differences in arborization volumes throughout the central nervous system (Cachero *et al*. 2010; Yu *et al*. 2010), which would likely be directed by differences in cell adhesion/connectivity properties of the neurons. This led to predictions that differences in neuronal connectivity are important mechanisms to mediate behavioral dimorphism (Cachero *et al*. 2010; Yu *et al*. 2010).

In-depth *in vitro* analyses of protein-protein interactions have shown that each Dpr has dimeric interactions with specific DIP proteins, with some having multiple DIP interacting partners. Additionally, some Dprs interact dimerically with Dprs through either heterophilic or homophilic interactions, and some of the DIPs interact dimerically through homophilic interactions (Ozkan *et al*. 2013; Carrillo *et al*. 2015; Cosmanescu *et al*. 2018)(summarized in **Supplemental Table 1**). Functional analyses of the Dprs and DIPs have revealed roles in synaptic connectivity and specificity of neuronal targeting in the *Drosophila* neuromuscular junction, visual system and olfactory system (Carrillo *et al*. 2015; Tan *et al*. 2015; Barish *et al*. 2018; Xu *et al*. 2018; Ashley *et al*. 2019; Courgeon and Desplan 2019; Menon *et al*. 2019; Venkatasubramanian *et al*. 2019; Xu *et al*. 2019). Cell adhesion molecules have already been shown to be important for sculpting dimorphism in *fru P1* neurons, with studies of the IgSF member encoded by *roundabout* (*robo*) shown to be a direct target of Fru^M^ and responsible for dimorphic projections and morphology (Mellert *et al*. 2010; Ito *et al*. 2016). Thus, the Dprs/DIPs are good candidates for directing sexual dimorphism in connectivity and morphology that underlies differences in reproductive behavior.

Our inroad into the study of the role of Dprs/DIPs in *fru P1* neurons came from a systematic reanalysis of several genomic studies that shows that all the *dprs* and *DIPs* examined are potentially regulated by Fru^M^ or are expressed in *fru P1* neurons. Additionally, a live tissue, *in vivo* staining approach demonstrates that there is sexual dimorphism in the overlap of *fru P1* neurons that stain with a Dpr or DIP. This prompted us to examine the sets of neurons that express *fru P1* and one of the *dprs* or *DIPs*, using a genetic intersectional strategy (*fru P1* ∩ *dpr/DIP;* **Figure 1C**), to gain insight into the combinatorial codes of cell adhesion molecules that direct development of *fru P1*-expressing neurons in males and females. Additionally, we examine the roles of neurons expressing *fru P1* and a *dpr* or *DIP* in reproductive behaviors to gain insight into whether the *dprs/DIPs* expression repertoires provides insights into functions of neuronal subtypes in directing behavior. In addition, this allows us to begin to elucidate which combinations of neurons underlie discrete steps in the courtship ritual. Additional genetic perturbation screens reveal functional roles of the sex hierarchy, and *DIP-ε*, in establishing sex-specific architecture of *fru P1 ∩ DIP-α* neurons. A single cell RNA-sequencing analysis demonstrates the myriad, unique combinations of *dprs/DIPs* expressed in individual *fru P1* neurons, with overlapping expression of at least one *dpr* or *DIP* in every *fru P1* neuron examined. Additionally, these single cell analyses generally show that *dprs* are expressed in more neurons at intermediate levels, whereas *DIPs* have higher expression in fewer neurons. Taken together, the *dprs* and *DIPs* play critical roles in establishing the *fru P1* neural circuitry in both males and females.

## Results

### Genome-wide studies provide evidence that *dprs* and *DIPs* function in *fru P1*-expressing neurons

Our systematic reanalysis of previous genomic studies shows that *dprs* and *DIPs* likely have a role in *fru P1* neurons (**Figure 1B**), with the majority of the *dpr/DIP* genes in the analysis identified as regulated by Fru^M^ or expressed in *fru P1* neurons, in at least three independent genome-wide studies (Goldman and Arbeitman 2007; Dalton *et al*. 2013; Neville *et al*. 2014; Vernes 2014; Newell *et al*. 2016). Furthermore, a DNA binding site analysis further confirms this regulation. There is alternative splicing at the 3’ end of *fru P1* transcripts that results in one DNA-binding-domain-encoding-exon being retained out of five potential exons. The predominant isoforms of Fru^M^ contain either the A, B or C DNA binding domain in the central nervous system (binding sites and genome-wide analysis described in Dalton *et al*. 2013). When we search for the presence of the three sequence motifs near/in the *dpr/DIP* loci, Fru^M^ binding sites are found near/in all but two *dpr/DIP* loci that are examined (**Supplemental Table 1**). Therefore, a systematic reanalysis of genome-wide studies strongly supports a role of *dpr/DIPs* in *fru P1*-expressing neurons.

### Live tissue staining shows sexual dimorphism in the number of cells that overlap with Dpr/DIP binding and *fru P1* neurons

We perform live tissue, *in vivo* staining, using conditioned tissue culture media that contains the epitope-tagged, extracellular regions of a Dpr or DIP. This allows us to examine binding to their respective Dpr/DIP partners in brain tissues of 48-hour pupae and 0-24 hour adults (as done in Fox and Zinn 2005; Lee *et al*. 2009; Ozkan *et al*. 2013). Using this approach, we detect signal for two Dprs and two DIPs in the subesophageal ganglion of the brain (Dpr3, Dpr16, cDIP, and DIP-γ; **Supplemental Figure 1**). The live staining technique is not effective throughout the adult brain and for all Dprs/DIPs tested, perhaps due to the inability of the epitope-tagged Dprs/DIPs extracellular regions to penetrate other regions in live brain tissues, which are not permeabilized by detergent, as is done for fixed tissue. The number of neuronal cell bodies with staining is similar in males and females at both time points, in wild type and *fru P1* mutants, with some significant differences with small effect sizes. However, the number of neuronal cell bodies with staining that overlap with *fru P1* is significantly higher in males compared to females at both time points. Given that we do not see large sex-specific changes in the number of cells with signal in *fru P1* mutants, suggests that regulation of *dprs/DIPs* is more complex than simple regulation by *fru P1*. Overall, the analysis reveals sexual dimorphism in binding of tagged Dpr/DIP proteins to *fru P1* neurons in the subesophageal ganglion brain region using a live staining approach (**Supplemental Figure 1**), with more neurons with overlap detected in males.

### A genetic intersectional approach identifies neurons that express both *fru P1* and a *dpr* or *DIP* in males and females

The above results led us to examine the expression patterns of neurons that express both *fru P1* and a *dpr* or *DIP*, using a genetic intersectional approach (**Figure 1C**). This approach restricts expression of a membrane-bound-GFP marker to neurons with intersecting expression of *fru P1* and a *dpr* or *DIP (fru P1* ∩ *dpr/DIP)*. This is accomplished using a UAS-membrane-bound GFP reporter transgene that requires removal of an FRT-flanked stop cassette for expression. Removal of the stop cassette is mediated by *fru P1* driven FLP recombinase (Yu *et al*. 2010). This system is used in combination with a collection of *dpr*- and *DIP-Gal4* transgenic strains (**Figure 1C**) (Venken *et al*. 2011; Nagarkar-Jaiswal *et al*. 2015a; Nagarkar-Jaiswal *et al*. 2015b; Tan *et al*. 2015; Lee *et al*. 2018). We primarily focus the analysis on 4-7 day adults (**Figures 2 and 3**), which are sexually mature adults, and 0-24 hour adults to determine if the patterns change during early adult stages (**Supplemental Figures 2 and 3**). Additionally, behavioral studies are performed in 4-7 day adults (**Figures 4–6**), so the expression and behavioral data can be co-analyzed (**Figure 7**). At a gross morphological level, the patterns we observe in older 4-7 day old adults are also present in 0-24 hour adults, though in some cases expression in the mushroom was not as robust at the early time point.

**Figure 2.**
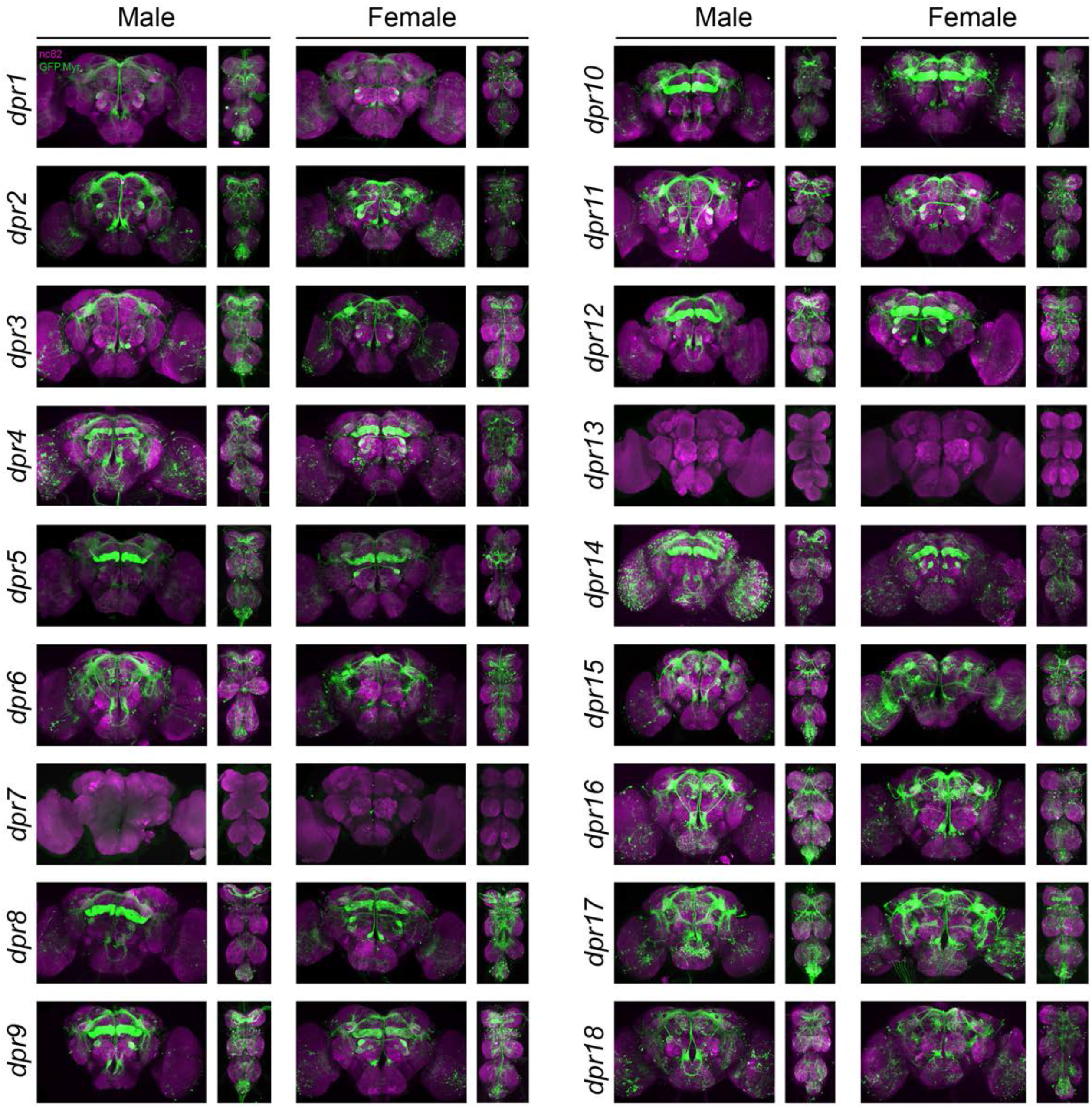
Visualization of *fru P1 ∩ dpr/DIP* neurons. Maximum intensity projections of brain and ventral nerve cord tissues from 4-7 days old male and female flies. The *fru P1∩ dpr/DIP* intersecting neurons are labeled with green (rabbit α-GFP Alexa Flour 488), and neuropil are labeled with magenta (mouse α-nc82, Alexa Flour 633). The genotype is *dpr/DIP-Gal4/10xUAS > stop > GFP.Myr; fru P1^FLP^*, except for *dpr4, dpr14, dpr18, dpr19* and *DIP-ι*. These five *Gal4* transgenic strains were generated using a CRISPR mediated insertion of the *T2A-Gal4* with the dominant 3xP3-GFP marker. For this strain, *10xUAS > stop > myr::smGdP-cMyc* was used and *fru P1∩ dpr/DIP* intersecting neurons are labeled with red (rabbit α-Myc, Alexa Flour 568) and then false-colored to green. The neuropil are labeled with magenta (mouse α-nc82, Alexa Flour 633). Four Gal4s did not show expression upon intersecting: *dpr7, dpr13, dpr19*, and *DIP-iota. dpr7* and *dpr13* have expression with 10xUASmCD8gfp confirming the Gal4s can drive expression outside of *fru P1*-expressing neurons. *dpr19* and *DIP-iota* were tested with 10xUAS-RFP, and only *DIP-iota* showed expression outside of *fru P1*-expressing neurons.

**Figure 3.**
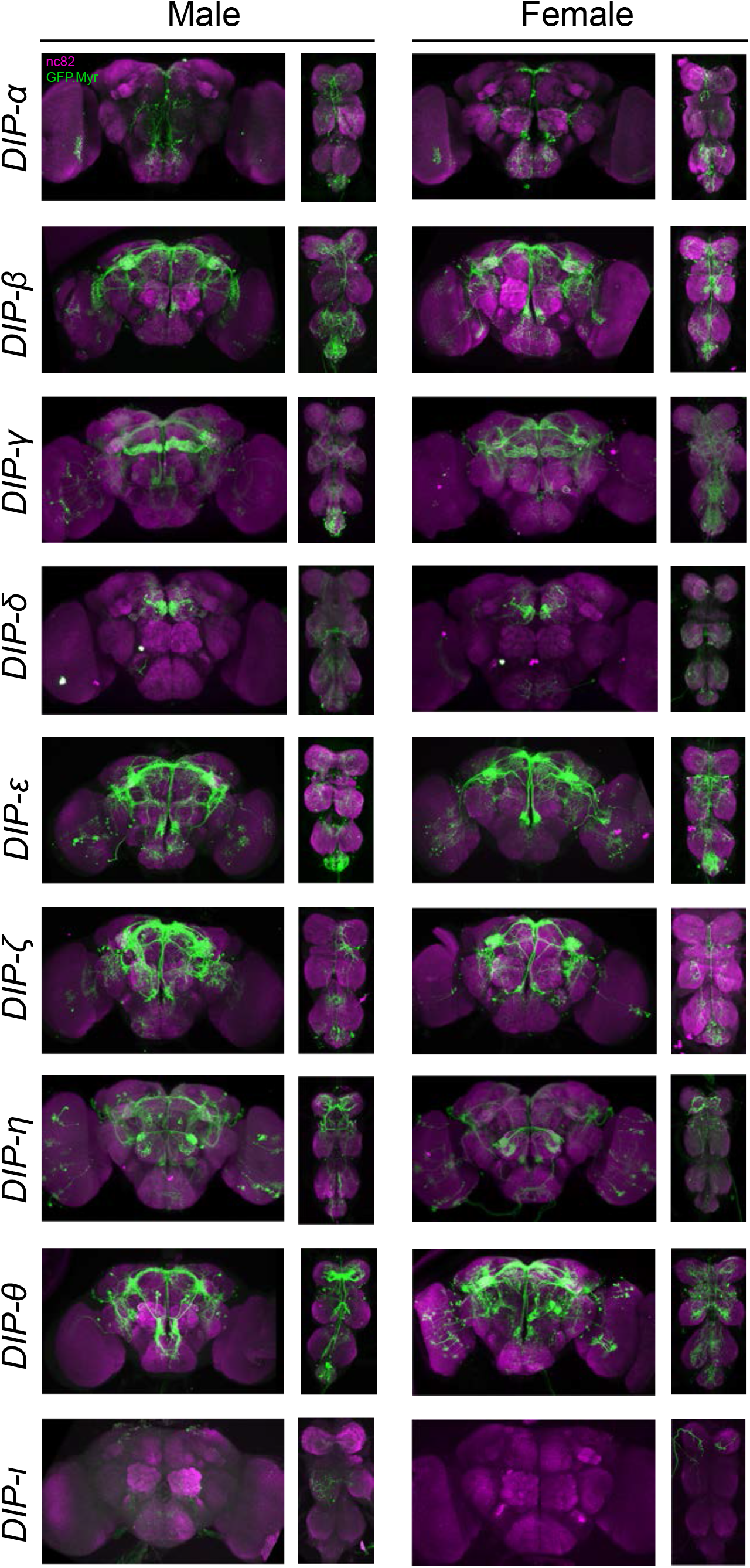
Visualization of *fru P1 ∩ dpr/DIP* neurons. Maximum intensity projections of brain and ventral nerve cord tissues from 4-7 days old male and female flies. The *fru P1∩ dpr/DIP* intersecting neurons are labeled with green (rabbit α-GFP Alexa Flour 488), and neuropil are labeled with magenta (mouse α-nc82, Alexa Flour 633). The genotype is *dpr/DIP-Gal4/10xUAS > stop > GFP.Myr; fru P1^FLP^*, except for *dpr4, dpr14, dpr18, dpr19* and *DIP-ι*. These five *Gal4* transgenic strains were generated using a CRISPR mediated insertion of the *T2A-Gal4* with the dominant 3xP3-GFP marker. For this strain, *10xUAS > stop > myr::smGdP-cMyc* was used and *fru P1∩ dpr/DIP* intersecting neurons are labeled with red (rabbit α-Myc, Alexa Flour 568) and then false-colored to green. The neuropil are labeled with magenta (mouse α-nc82, Alexa Flour 633). Four Gal4s did not show expression upon intersecting: *dpr7, dpr13, dpr19*, and *DIP-iota. dpr7* and *dpr13* have expression with 10xUASmCD8gfp confirming the Gal4s can drive expression outside of *fru P1*-expressing neurons. *dpr19* and *DIP-iota* were tested with 10xUAS-RFP, and only *DIP-iota* showed expression outside of *fru P1*-expressing neurons.

**Figure 4.**
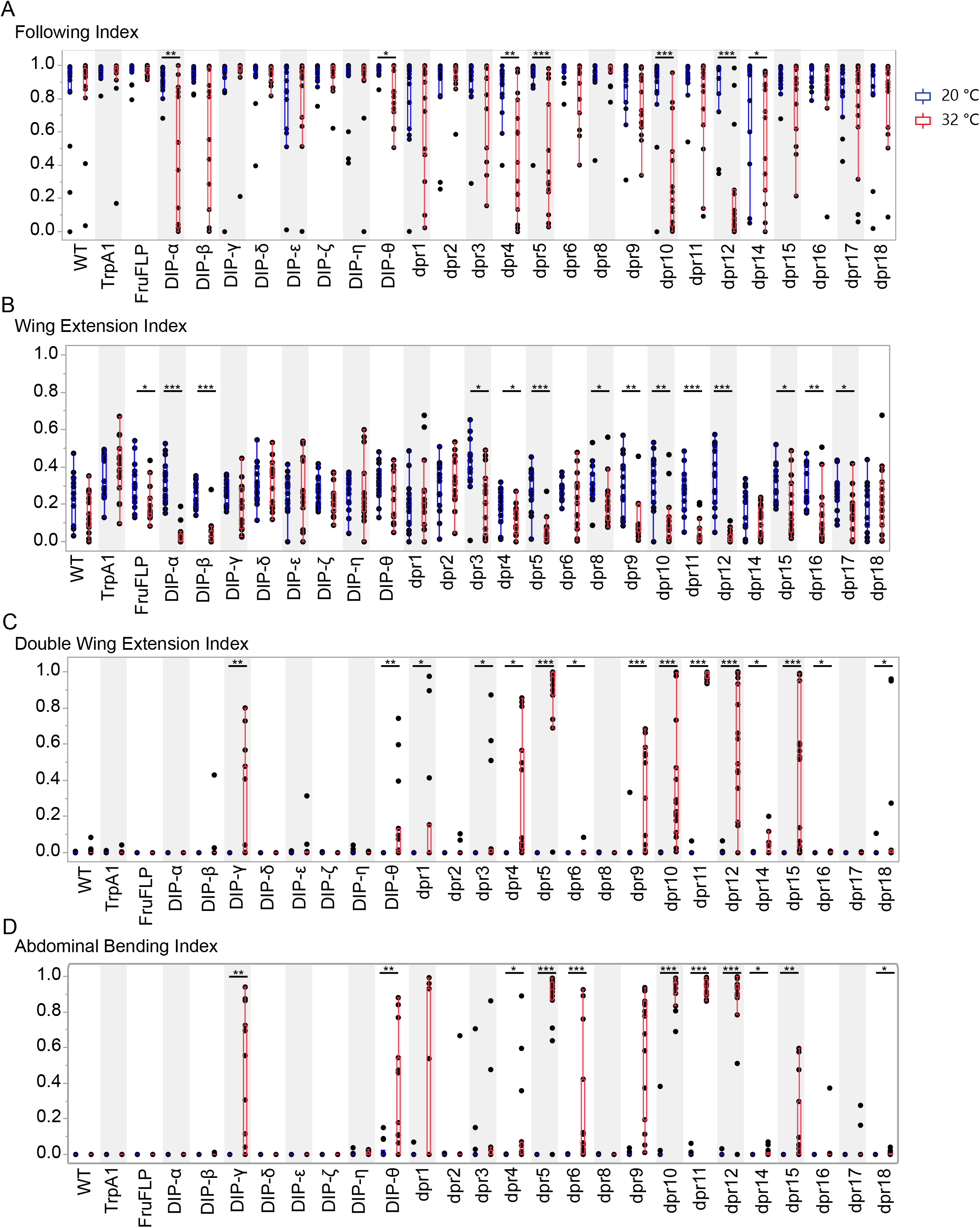
Activation of *fru P1* ∩ *dpr/DIP* intersecting neurons results in atypical courtship behaviors. Courtship behaviors of *dpr/DIP-Gal4/ UAS > stop > TrpA1; fru P1^FLP^* males were recorded at the control temperature (20°C, blue box plots) and the activating temperature for TrpA1 (32°C, red box plots). The control genotypes are the wild type strain Canton S, and the *UAS > stop > TrpA1* and *fru P1^FLP^* single transgenes, which were crossed to Canton S. Virgin Canton S (*white*) females were used as targets. (**A**) Following index is the fraction of time the male spent oriented towards or chasing the female around the chamber. (**B**) Wing extension index is the fraction of time the male spent unilaterally extending and vibrating his wing. (**C**) Double wing extension index is the fraction of time the male spent extending and vibrating both wings simultaneously. (**D**) Abdominal bending index is the fraction of time the male spent curling his abdomen under. The lines on the quantile box plot correspond to the quantiles in the distribution output, with the center line as the median. The whiskers extend from the 1^st^ and 3^rd^ quartiles to the edges, which correspond to the minimum and maximum values, excluding outliers. The nonparametric Wilcoxon rank sum test was used to test for significant difference between control and activating temperature within each genotype. n=15 * < 0.05 **<0.005 ***<0.0005. All lines were examined for expression of TrpA1 to confirm the system was working effectively (data not shown).

**Figure 5.**
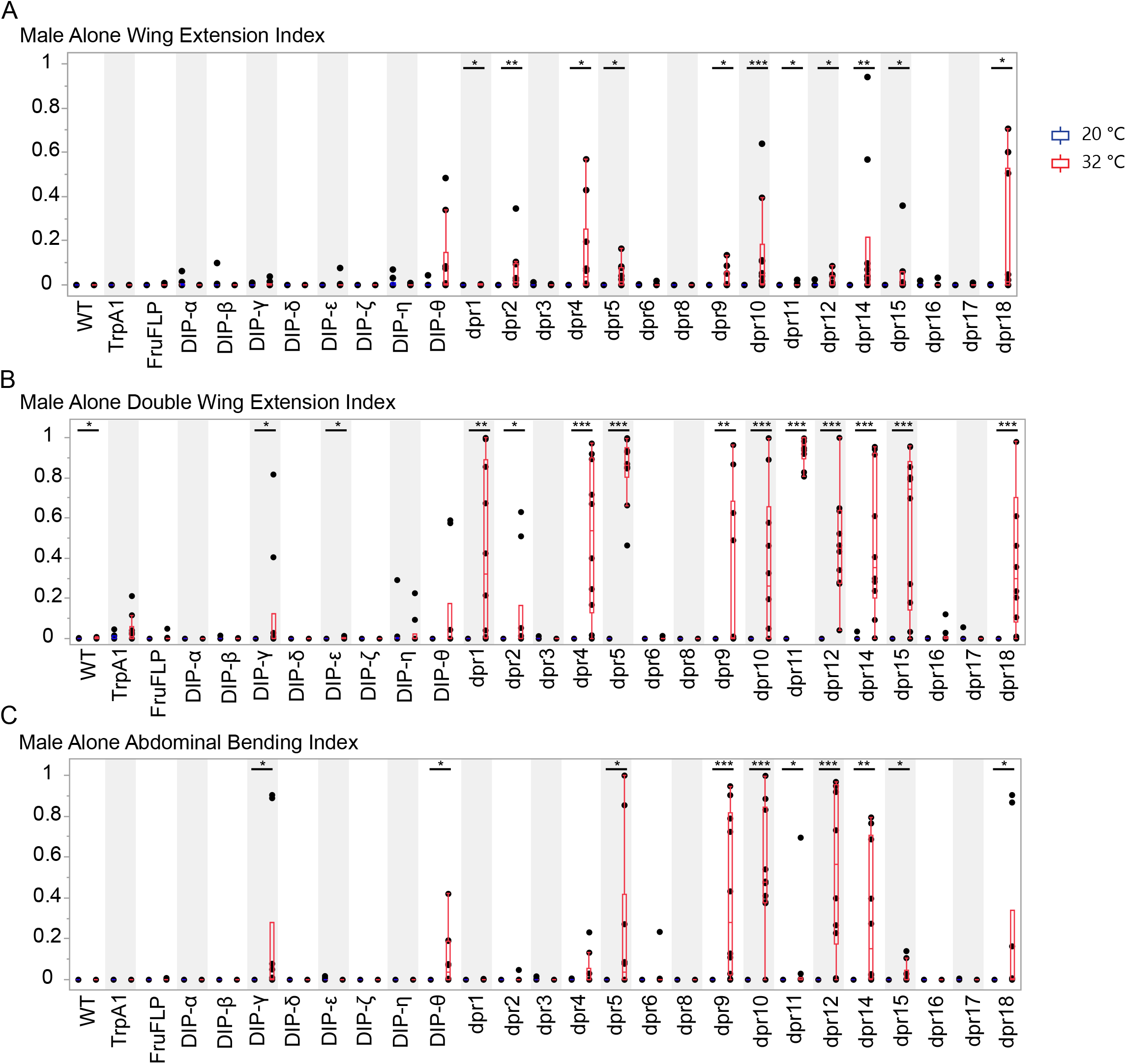
Activation of *fru P1* ∩ *dpr/DIP* intersecting neurons is sufficient to induce courtship behaviors in solitary males. Courtship behaviors of *dpr/DIP-Gal4/ UAS > stop > TrpA1; fru P1^FLP^* solitary males were recorded at the control temperature (20°C, blue box plots) and the activating temperature (32°C, red box plots). The control genotypes are the wild type strain Canton S, and the *UAS > stop > TrpA1* and *fru P1^FLP^* single transgenes, which were crossed to Canton S. (**A**) Wing extension index, (**B**) Double wing extension index (**C**) Abdominal bending index, and quantile box plots are as described in Figure 3. The nonparametric Wilcoxon rank sum test was used to test for significant difference between control and activating temperature within each genotype. n=10 * *P*< 0.05 \*\**P*<0.005 \*\*\**P*<0.0005.

**Figure 6.**
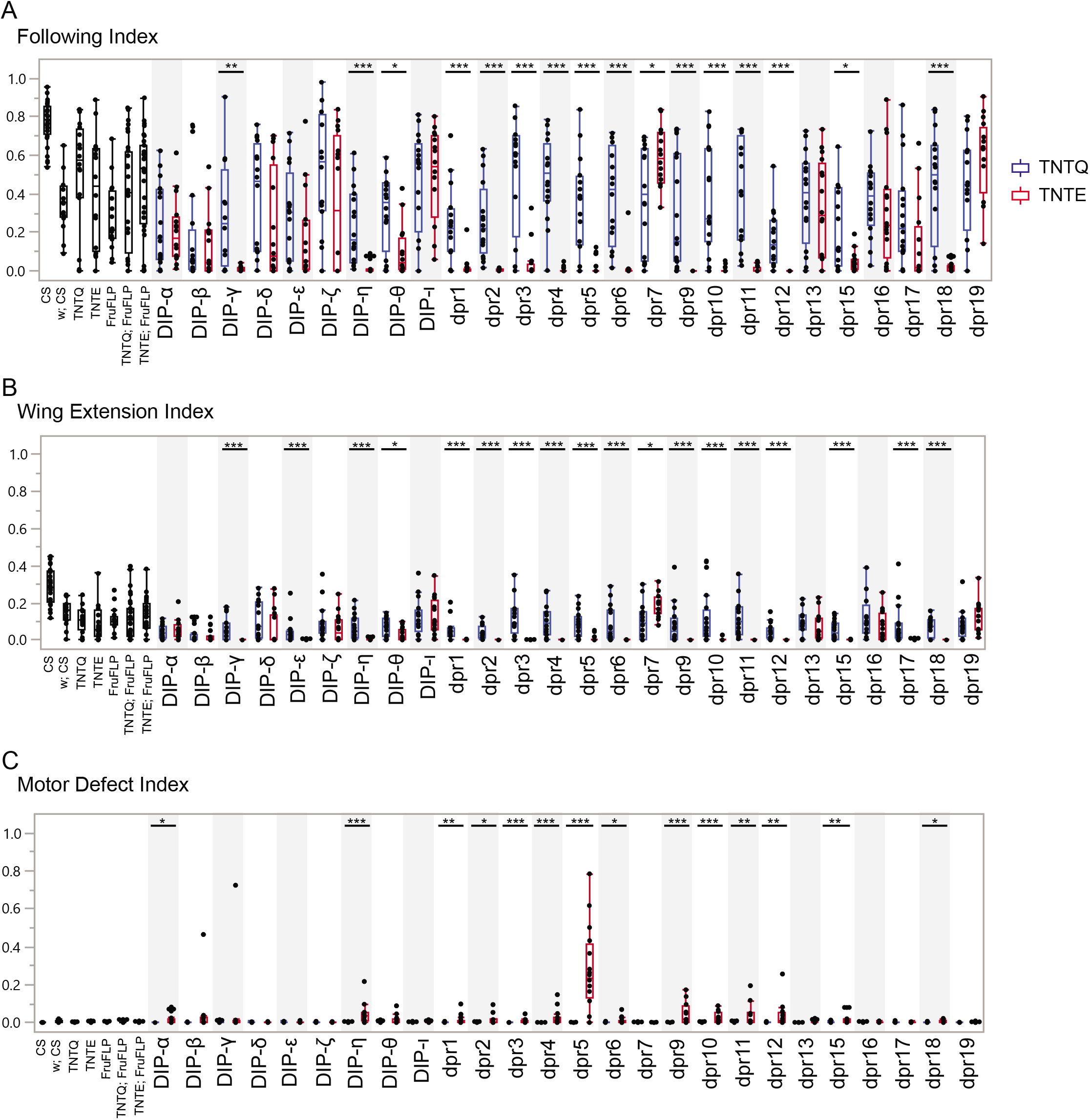
Silencing *fru P1* ∩ *dpr/DIP* intersecting neurons results in atypical courtship and severe motor defects. Courtship behaviors of *dpr/DIP-Gal4/ UAS > stop > TNTQ; fru P1^FLP^* (control condition, blue boxplots) and of *dpr/DIP-Gal4/ UAS > stop > TNTE; fru P1^FLP^* (experimental condition, red boxplots) males were quantified. Control genotypes (black boxplots) are the wild type strain Canton S and Canton S (*white*), *fru P1^FLP^ UAS > stop > TNTQ*, and *UAS > stop > TNTE* single transgenes, as well as *UAS > stop > TNTQ; fru P1^FLP^* and *UAS > stop > TNTE; fru P1^FLP^* double transgenes. The single and double transgene controls were crossed to Canton S (*white*). The *dpr*- or *DIP-Gal4* is listed on the x-axis and the fraction of time spent performing the behavior is on the y-axis. (**A**) Following index, (**B**) wing extension index, and the quantile box plots are as described in Figure 4. (**C**) Motor defect index is the fraction of time the fly spent on his back after falling. The nonparametric Wilcoxon rank sum test was performed to determine significant differences between experimental and control conditions with the same *dpr/DIP-Gal4*. n=16 for all genotypes except for Canton S, and the double transgene controls, which have n=32. Those three genotypes were assayed twice, n=16 each time, to ensure consistency throughout the duration of the experiment and pooled for this analysis. The *dpr19-Gal4* did not produce an expression pattern in the nervous system, using both a *10XUAS-RFP* reporter and the intersectional approach, at the time points examined. n=16. \**P*<0.05 \*\**P*<0.005 \*\*\**P*<0.0005.

**Figure 7.**
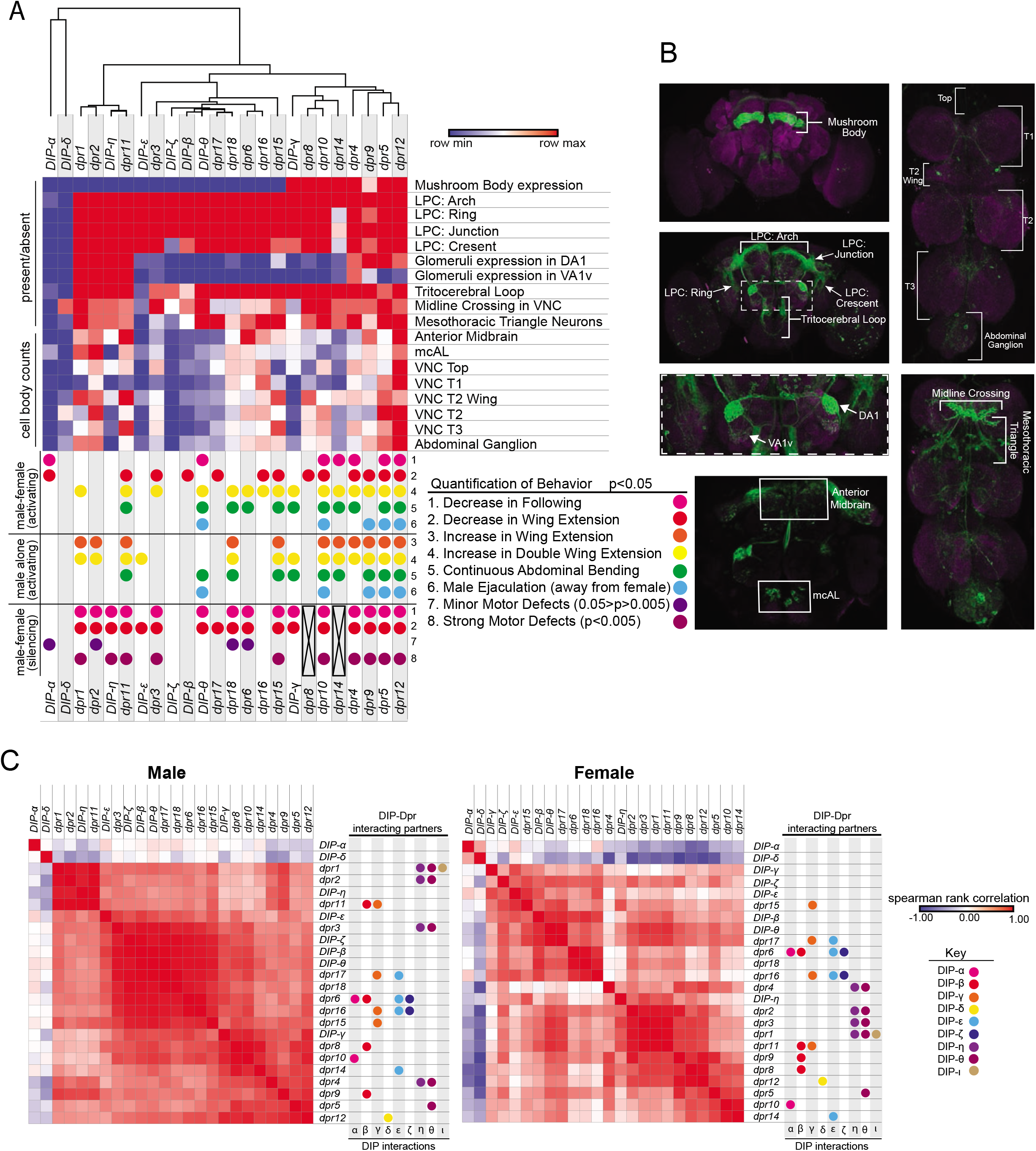
Meta-analysis of expression patterns of *fru P1* ∩ *dpr/DIP* intersecting neurons and behavior data. Meta-analysis using behavior data and image analysis data of 4-7-day old flies. (**A**) Heatmap of *fru P1* ∩ *dpr/DIP* intersecting neurons expression patterns in the male adult CNS. For each row, the minimum (blue), middle (white) and maximum (red) values are indicated. The top of the heatmap shows the relationship across the expression patterns of the *dprs* and *DIPs*, with a dendrogram. The summary of phenotypic analyses of male sexual behaviors, using either activating or silencing effector genes (see **Figures 4–6**), is shown below the heat map. The dot indicates a significant change in behavior (p<0.05, unless indicated). The black X indicates that there was no experimental progeny from the cross, due to lethality, and therefore were not tested behaviorally. (**B**) Labeled confocal images showing the morphological featured scored. (**C**) Correlation analysis of GFP expression results (male on left and female on right). The scale for the spearman correlation is −1 (blue) to 1 (red). The dots to the right indicate the DIP interacting partners for each Dpr (left-hand side of each graph) (Dpr-DIP interactome based on Carrillo *et al*. 2015). The full data set is provided (**Supplemental Table 2**)

Based on our examination of the expression patterns in 27 intersecting genotypes, we find that 24 showed clear, membrane-bound GFP expression in the central nervous system at the time points examined. Of these, only two *fru P1* ∩ *DIP* genotypes have very restricted and unique patterns (*fru P1* ∩ *DIP*-δ and *fru P1* ∩ *DIP-a*), whereas the other genotypes have broader expression, with many in similar regions/patterns (**Figures 2 and 3**). For example, 22 intersecting genotypes, in both males and females, have consistent expression in the brain lateral protocerebral complex, including within the arch, ring, junction and crescent (for summary see **Figure 7 and Supplemental Table 2**). This region has been shown to have *fru P1* neurons with sexually dimorphic arbor volumes (Cachero *et al*. 2010; Yu *et al*. 2010). Furthermore, the lateral protocerebral complex has inputs from sensory neurons and is predicted to be a site of sensory integration, to direct motor output (Yu *et al*. 2010). We find 8 intersecting genotypes have expression in mushroom bodies in both males and females. This region has a well-established role in learning and memory, including learning in the context of courtship rejection (McBride *et al*. 1999; Montague and Baker 2016; Jones *et al*. 2018; Zhao *et al*. 2018). Overall, the majority of *fru P1* ∩ *dpr/DIP* genotypes are expressed in similar regions, suggesting that some may function in combinatorial manner within a neuron to direct patterning and/or synaptic targeting.

We observe sex differences in the presence of morphological features and cell body number in regions we scored (**Figures 2 and 3 and Supplemental Table 2**), which were largely chosen because they were previously reported to display sexual dimorphism (Cachero *et al*. 2010; Yu *et al*. 2010). For example, 18 intersecting genotypes show consistent presence of signal in the mesothoracic triangle neuronal projections in males, but only two lines do so in females. While both males and female have expression in antennal lobe glomeruli DA1 and VA1v in several intersecting genotypes, there is also sexual dimorphism, with four genotypes having consistent expression in only female DA1 glomeruli (*fru P1* ∩ *dpr3, dpr10, dpr17, DIP-θ)*. In the ventral nerve cord, a midline crossing phenotype is consistently observed for the majority of intersecting genotypes only in males, which was previously shown to be a male-specific phenotype for a set of gustatory neurons (Mellert *et al*. 2010). For all regions where cell bodies are counted, the trend was that there are more cell bodies in males than females. Thus, the differences in the patterns of expression between males and females are not large, with several genotypes having quantitative differences in the numbers of cell bodies present, rather than a more complete presence or absence difference. It is possible that there are additional quantitative differences that are not detected based on the resolution of the analyses, including quantitative differences in expression level of *dpr/DIPs*, or their sub-cellular localization, or in regions/features that are not quantified here (**Figures 2 and 3 and Supplemental Table 2**).

### Activation of *fru P1 ∩ dpr/DIP* neurons results in atypical courtship behaviors

Substantial progress has been made in showing *fru P1* has a critical role in reproductive behaviors, including determining the function of small subsets of neurons that are responsible for different aspects of behavior (reviewed in Auer and Benton 2016). The tools in hand can further address if additional combinations or quantitative differences in the number of *fru P1* neurons are important for behavioral outcomes, given the *fru P1* ∩ *dpr/DIP* subsets and combinations we examine are distinct from those previously studied. We use the genetic intersectional strategy to activate intersecting neurons, by driving expression of TrpA1, a heat activated cation channel (**Figure 1C**) (von Philipsborn *et al*. 2011). This allows for temporal control of neuronal activation by an acute increase of the temperature in the courtship chambers (32°C; controls were at 20°C). We find that neuronal activation resulted in decreases in male following and wing extending towards females for several genotypes (**Figure 4 and 7 and Supplemental Table 3**). We also observe that neuronal activation of *fru P1* ∩ *dpr* (13/16) and *fru P1* ∩ *DIP* (2/8) genotypes caused atypical courtship behavior towards a female, including double wing extension, and continuous abdominal bending, even if the female had moved away (**Figure 4 and 7**). These atypical behaviors could account for some of the decreases in following and wing extension. For example, if a male is locked into abdominal bending, this would reduce courtship following behavior. Additionally, we find that some males ejaculated on the chamber in five intersecting genotypes: *dpr5* (5 /15), *dpr9* (3 /15), *dpr10* (3 /15), and *dpr12* (2 /15), and *DIP-θ* (4 /15). Of note, *fru P1* ∩ *DIP-α* is the only strain that showed a decrease in courtship activities without a concomitant increase in atypical courtship behaviors. This suggests that *fru P1* ∩ *DIP-α* neurons may normally inhibit courtship behaviors when they are activated.

We next determine if the males require females to reach an arousal threshold needed to perform typical and atypical courtship behaviors, given that several of the courtship behaviors described above occur when the male was not oriented towards the female. To address this question, we examine courtship behaviors in solitary males, using the same temporal activation strategy as above. We find that activation of the *fru P1* ∩ *dpr/DIP* neurons is sufficient to elicit single wing extension, double wing extension, and abdominal bending in *fru P1* ∩ *dprs* (11/16) and *fru P1* ∩ *DIPs* (3/8) (**Figure 5, 7 and Supplemental Table 3**). Similarly, activating the intersecting *fru P1* neuronal populations of *fru P1* ∩ *dpr5* (5 /10), *dpr9* (1/10), *dpr10* (1/10), *dpr12* (3/10), and *DIP-θ* (1/10) causes males to ejaculate without a female present. Overall, activation of these subsets of *fru P1* neurons is sufficient to direct reproductive behaviors, even if a female is not present, consistent with other neuronal activation experiments (reviewed in Auer and Benton 2016).

### Silencing *fru P1 ∩ dpr/DIP* neurons result in courtship changes

Given that activation of *fru P1* ∩ *dpr/DIP* neuronal subsets resulted in changes in courtship behaviors, we next determine how silencing these neurons impacts male-female courtship, to gain further insight into their roles. To test this we use the genetic intersectional approach with a *UAS > stop > TNT* transgene (**Figure 1C**) (Stockinger *et al*. 2005). The intersecting genotypes express tetanus toxin light chain, which cleaves synaptobrevin, resulting in synaptic inhibition (Sweeney *et al*. 1995). As a control we also examine courtship behaviors of flies expressing an inactive form of *TNT* (TNTQ), using the genetic intersectional approach.

In addition to scoring courtship behaviors, motor impairment is also scored (**Figure 6 and Supplemental Table 3**). Given that neuronal silencing in several genotypes results in motor impairment, in which the male fell and is unable to quickly right himself, we quantify the time when the fly could not right himself as “motor defect” and subtract this from the overall courtship time for behavioral indices (**Figure 6**). The intersecting genotypes that consistently demonstrate motor defects additionally show decreases in following and wing extension upon silencing, likely due to some motor impairment (*fru P1* ∩ *dpr1, dpr3, dpr4, dpr5, dpr9, dpr10, dpr11, dpr12, dpr15* and *DIP-η)*. Additional courtship behavioral indices and latencies are quantified and those with motor defects show additional strong courtship phenotypes (**Supplemental Table 3**). However, seven intersecting genotypes have a decrease in following/wing extension indices and only minor or no motor impairment (*fru P1* ∩ *dpr2, dpr6, dpr17, dpr18, DIP-ε, DIP-θ, andDIP-γ)*. One genotype, *fru P1* ∩ *dpr7*, has an increase in following/wing extension with neuronal silencing. In the case of *fru P1* ∩ *dpr7*, we do not detect GFP expression in the central or peripheral nervous system in adults, so the neurons underlying this phenotype remain to be determined. Locomotor activity of the seven intersecting genotypes with no or minor motor defects are further analyzed for motor impairment (p<0.005 for strong motor defects; 0.05>p>0.005 for minor; **Supplemental Table 3**), along with *fru P1* ∩ *dpr7*, and *fru P1* ∩ *dpr10*, which has strong motor impairment. If there is a significant difference, the intersecting genotype with neuronal silencing has increased locomotor activity in the activity monitors, suggesting that the courtship phenotypes are not due to overall loss in motor activity (**Supplemental Table 3**).

As above in the neuronal activating experiments, silencing *fru P1* ∩ *Dprs* (13/19) is more likely to cause a courtship defect than silencing *fru P1* ∩ *DIPs* (4/9). Given the large effect size of the courtship defects compared to the smaller effect size of the motor defect, it is clear that silencing *fru P1* ∩ *dpr/DIP* neurons in the central nervous system, for most genotypes, suppresses courtship (**Figure 6**). This is consistent with previous studies that have found that silencing *fru P1* neurons in males leads to decreased courtship towards a female (Manoli *et al*. 2005; Stockinger *et al*. 2005). Interestingly, *fru P1* ∩ *DIP-α* is the only strain to demonstrate motor defects, but no change in courtship behaviors upon silencing, underscoring the previous hypothesis that these neurons may normally be inhibitory for courtship.

### Meta-analysis of male *fru P1* ∩ *dpr/DIP* expression patterns and behavioral data

Next, we determine if intersecting genotypes with similar expression patterns also have similar behavioral outcomes in the neuronal activating and silencing experiments described above. We use a heuristic approach and generate a heatmap that groups *dprs/DIPs* based on similarity of the *fru P1* ∩ *dpr/DIP* membrane-bound-GFP expression data (**Figure 7A, and Supplemental Table 2** for additional visualizations). At the top of the heatmap is a dendrogram showing the relationships in expression data, grouping those that are most similar together (from data in **Figure 2 and 3 and Supplemental Table 2**). The bottom has colored dots that indicate the behavioral changes observed in the three different behavioral perturbation data sets (from data in **Figures 4–6**). The scoring key for the GFP expression phenotypes is shown (**Figure 7B and Supplemental Table 2**). Only the 24 intersecting genotypes with GFP expression data are included in the heat map.

There is a set of eight intersecting genotypes grouped together on the right of the dendrogram that all have expression in the mushroom body and several regions within the lateral protocerebral complex, but varied expression across the other morphological features (**Figure 7A**; *fru P1* ∩ *dpr4, dpr5, dpr8, dpr9, dpr10, dpr12, dpr14* and *DIP-γ*). Seven have similar types of atypical courtship behaviors in the activating experiments (excluding *fru P1* ∩ *dpr8*), in the male-female courtship assays. These seven also have similar behavioral phenotypes in the male-alone condition, indicating that the activation threshold in these lines can be achieved without a female present (**Figure 5**).

Furthermore, among the eight genotypes, there are four intersecting genotypes that have male ejaculates in the chamber, in both the male-female and male-alone neuronal activation assays. All four intersecting genotypes also have relatively high cell body counts in the abdominal ganglion, a region in the ventral nerve cord that has previously been shown to drive ejaculation (**Supplemental Table 2**) (Tayler *et al*. 2012). However, not all intersecting genotypes with expression in the abdominal ganglion show the ejaculation phenotype, as shown in the heatmap. Furthermore, there is an intersecting genotype that does not have mushroom body expression, but also has the ejaculation phenotype (*fru P1 ∩ DIP-θ*). These results reveal how different combinations and numbers of neurons can direct a similar behavioral outcome. Overall, the results point to a critical role for interactions between the mushroom body and protocerebral complex in directing courtship behaviors, which are modified by being activated in combination with other neuronal populations. This is consistent with an idea put forth previously that posited connections between these two brain regions may integrate diverse external stimuli with internal physiological state and previous behavioral experience (Yu *et al*. 2010).

Twenty-two intersecting genotypes have expression in different regions of lateral protocerebral complex, but no consistent expression in the mushroom body. An examination of the behavioral phenotypes reveals no consistent behavioral phenotypes, based on the lateral protocerebral complex expression data. While the lateral protocerebral complex is critical for higher order processing, the data further supports the idea that interactions across different combinations of activated neurons, in each intersecting genotype, is critical for the behavioral outcomes and underscores how different patterns of neuronal activity can direct similar behavioral outcomes.

### Correlation of *fru P1* ∩ *Dpr/DIP* expression patterns

As an additional heuristic tool, we plot the correlation of the GFP expression patterns for the male and female data (**Figure 7C and Supplemental Table 2**). One goal is to gain insight into whether Dprs/DIPs with the same interacting partners are co-expressed together. This allows us to gain insight into the mechanisms used by these IgSF molecules to direct cell adhesion and to determine if there are sex differences. Another goal is to determine if the protein-protein interactions may occur through *cis* (within the same neuron) vs *trans* (across neurons) interactions. For example, if protein-protein interactions are in *cis*, then the Dpr/DIP interacting partners will be expressed in the same neurons and have correlated expression patterns. To address these questions, the plots are annotated with DIPs (colored dots) that each Dpr interacts with on the right (based on interactome from Carrillo *et al*. 2015).

It appears that some Dprs/DIPs that bind the same partner have the most similar expression patterns. For example, in males *fru P1* ∩ *dpr1*, and *dpr2* have highly correlated expression and both Dpr1 and Dpr2 interact with DIP-η and DIP-θ. In addition, the male *fru P1* ∩ *DIP-η* expression pattern is highly correlated with *fru P1* ∩ *dpr1*, and *dpr2*, suggesting that Dpr-DIP protein-protein interactions may also occur in *cis*. Similarly, in females, *fru P1* ∩ *dpr1, dpr2*, and *dpr3* have highly correlated expression, with Dpr1, Dpr2 and Dpr3 also all interacting with DIP-η and DIP-θ. On the other hand, in males, *dpr11* does not have highly correlated expression with *DIP-β* and *DIP-γ*, though Dpr11 interacts with DIP-β and DIP-γ. This is consistent with protein-protein interactions occurring in *trans*. In females, *fru P1* ∩ *dpr8, dpr9, dpr11* (interact with DIP-β and DIP-γ) have highly correlated expression patterns, which is not observed in males. Therefore, there are sex-differences in the co-expression patterns of Dprs that could underlie dimorphism in morphology. *fru P1* ∩ *DIP-α* and *DIP-δ* have the most restricted expression patterns and they are not highly correlated with the expression patterns of their interacting Dpr partners, in either males or females. Overall, based on the correlation patterns in the expression data, it appears that some protein-protein interactions can occur in *cis* or *trans*. Additionally, some Dpr and DIPs with similar binding partners have correlated expression, which could be a mechanism to mediate the strength of neuronal adhesion. These observations are also supported by the single cell sequencing data (see below).

### A higher resolution analysis of *fru P1 ∩ DIP-α* reveals additional sexually dimorphic expression patterns

To gain insight into mechanisms that generate sexual dimorphism in morphology, we examine the relatively small number of *fru P1 ∩ DIP-α* neurons in male and females. Their small number facilitates in-depth analysis, as cell bodies and projection patterns are easier to discern (**Figure 8 and Supplemental Table 4**). While the overall patterns are similar (**Figure 8A**), there are fine-scale differences (**Supplemental Table 4**). There are sex-differences in the superior medial protocerebrum region (SMP; **Figure 8A and B, subpanels I**), where females have a longer (dotted-line) and broader projection (arrowhead), as compared to males. Moreover, in the medial part of midbrain, an “M” shaped peak forms (“M”-like) in males that is not typically observed in females (curved dotted-line, **Figure 8A and B, subpanels II&III**). Additionally, in the ventral lateral protocerebrum region (VLP) there are neuronal cell bodies (arrowhead, **Figure 8A and B, subpanels II&III**), and projections in a “square” shaped pattern that are more frequently observed in females (closed dotted-line, **Figure 8A and B, subpanels II&III**). There is also a greater frequency of neuronal cell bodies present in the subesophageal ganglion (SEG) in females, as compared to males (arrowhead, **Figure 8A and B, subpanels IV**). In the abdominal ganglia (AbG) of the ventral nerve cord there is a higher density of projections in males (**Figure 8A and B, subpanels V**). In contrast, females have a distinct “forceps” shaped pattern in the AbG region (arrowhead, **Figure 8A and B, subpanels V**). Taken together, it appears that the sex differences are due to differences in the number of neurons and also in the morphology of projections and arborizations (**Figure 8**).

**Figure 8.**
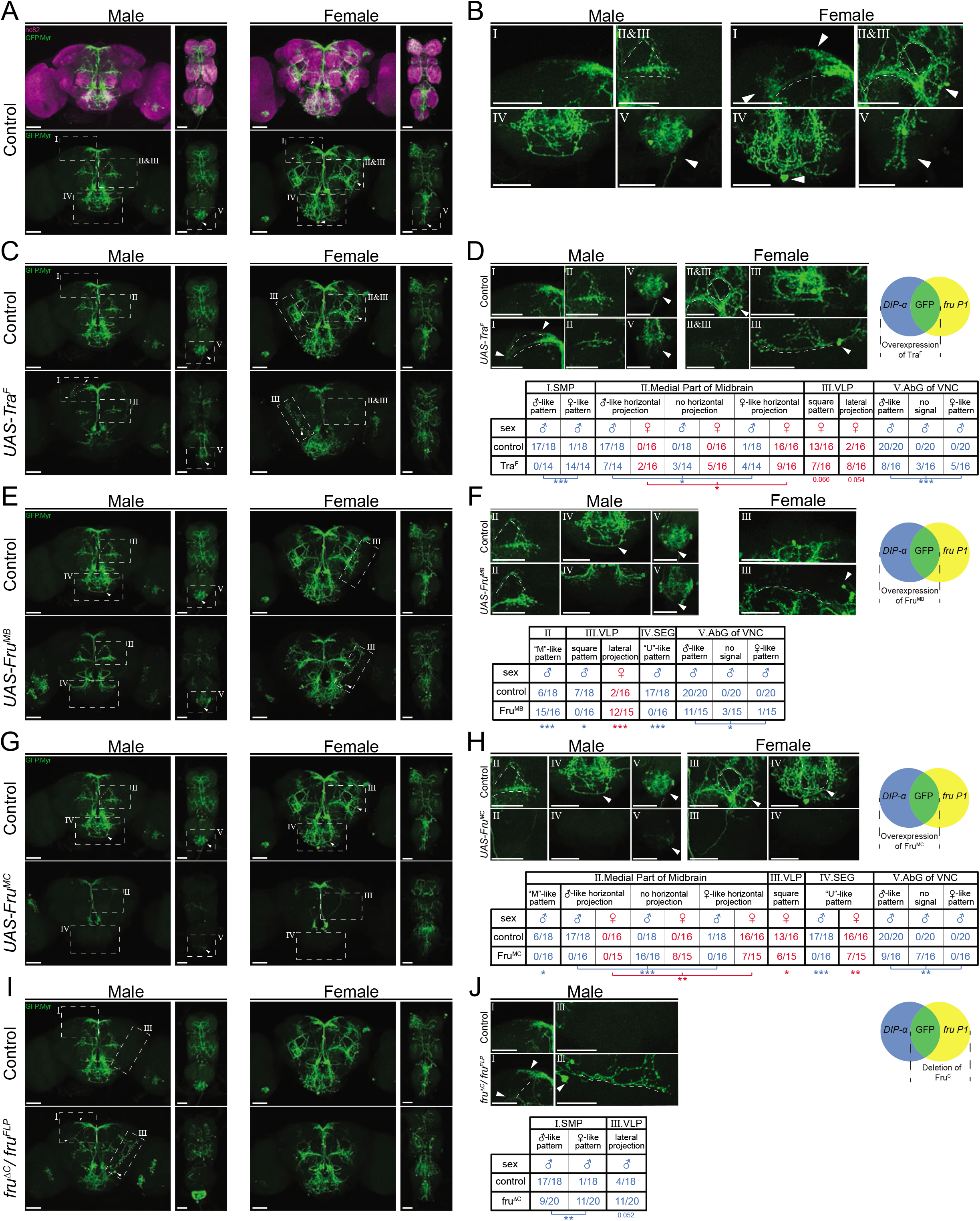
Higher resolution analyses of *fru P1* ∩ *DIP-α* neurons with sex hierarchy perturbations. Confocal maximum intensity projections of brains and ventral nerve cords from 4-7-day old adult flies. *fru P1* **∩** *DIP-α* neurons are in green (rabbit α-GFP Alexa Flour 488). Staining with the α-nc82 neuropil maker shows brain morphology in magenta (mouse α-nc82, goat α-mouse Alexa Flour 633). Image data were captured with 20x objective, with scale bars showing 50 μM (**A-J**). Higher magnification images were generated using the Zeiss Zen software package (**B, D, F, H and J**). Roman numerals are consistent across the panels in the same row. Venn diagrams show where membrane-bound GFP and sex hierarchy transgenes are expressed. **(A)** *fru P1* **∩** *DIP-α* expression patterns in males and females. **(B)** Computationally magnified images, with sexually dimorphic regions indicated. Subpanels show: **[I]** superior medial protocerebrum (SMP) region of the brain; **[II and III]** medial part of midbrain region, where there are horizontal projections, and the “M”-like pattern (more frequent in males). The square pattern (more frequent in females) is in the ventral lateral protocerebrum (VLP) region of the brain. The medial horizontal projection is in a more exterior section of the confocal stack then the other features **[II and III**]; **[IV]** Subesophageal ganglion region of the brain (SEG). The U-like pattern and a set of cell bodies more frequently found in females are shown; **[V]** The abdominal ganglion of the ventral nerve cord (AbG). **(C-J)** Examination of morphology of *fru P1* **∩** *DIP-α* neurons when sex hierarchy transgenes are expressed in *DIP-α* neurons. The quantification and statistics are provided in a table within the subpanel on the right of each row. This figure only shows regions that had significant changes due to sex hierarchy perturbation (full dataset provided; **Supplemental Table 4**). **(C-D)** Tra^F^ overexpression in *DIP-α* neurons. **[III]** a lateral projection in females that is not shown in wild type data in panel **B**. (**E-F)** Fru^MB^ overexpression in *DIP-α* neurons. **(G-H)** Fru^MC^ overexpression in *DIP-α* neurons, **(I-J)** Fru^C^ isoform deletion. Fru^MC^ is absent or highly reduced in *fru P1* neurons in this genotype, as transheterozygous for *fru^FLP^/fru^ΔC^*. Statistical significance of the differences in morphological features, between same sex control and genotypes with sex hierarchy transgene expression are indicated. Comparisons were done using the Fisher’s exact test (*P < 0.05, **P < 0.005, ***P < 0.0005). The morphological features with significant differences are indicated by lines below the table (male in blue and female in red). n ≥ 15 for each category. The genotypes of the samples shown are: *DIP-α^Gal4^; UAS>stop>GFP.Myr/+; fru^FLP^*/+ **(A-B)***, DIP-α^Gal4^; UAS>stop>GFP.Myr/ UAS-Tra^F^; fru^FLP^*/+ **(C-D)**, *DIP-α^Gal4^; UAS>stop>GFP.Myr/ UAS-Fru^MB^; fru^FLP^*/+ **(E-F)**, *DIP-α^Gal4^; UAS>stop>GFP.Myr/ UAS-Fru^MC^; fru^FLP^*/+ **(G-H),** *DIP-α^Gal4^; UAS>stop>GFPMyr/+; fru^FLP^/ fru^ΔC^* **(I-J)**. Brain region nomenclature are consistent with previous reports (Ito *et al*. 2014).

### Changing the sex of *DIP-α* neurons alters the *fru P1 ∩ DIP-α* co-expressing patterns

We next investigate whether perturbations of the sex hierarchy genes impact fine-scale sex differences in *fru P1 ∩ DIP-α* neurons (**Figure 8** and **Supplemental Table 4**). In this screen, *DIP-α-Gal4* drives broad expression of each transgene (see **Supplemental Table 4**), and the *fru P1 ∩ DIP-α* patterns are visualized. First, we examine the phenotypes when we overexpress the female-isoform of the sex hierarchy gene *tra* (*tra^F^*). This is expected to feminize the neurons by switching to female-specific splicing of *fru P1* (**Figure 1**). In males, the projections in the SMP became more female-like (**Figure 8C-D, subpanels I**). In the medial part of midbrain, the horizontal projections in half of the male samples are either more female-like or not detected (**Figure 8C-D, subpanels II**). Similarly, among half of the male samples, the neuronal patterns within the AbG are either more female-like or missing (**Figure 8C-D, subpanels V**). We observe unexpected phenotypes in females upon overexpressing Tra^F^, which suggests quantitative differences in Tra^F^ have biological outcomes, as we previously suggested (Arbeitman *et al*. 2016). For instance, a lateral ascending neuronal projection is observed more frequently in the VLP region (**Figure 8C-D, subpanels III** dotted line). However, the neuronal cell bodies in the VLP, the adjacent “square” shaped projection patterns (closed dotted-line, **Figure 8C-D, subpanels III**) and the medial horizontal projection (**Figure 8C-D, subpanels III**) are less frequently observed, as compared to control females.

We also examine phenotypes after Fru^M^ over-expression, by driving broad expression in *DIP-α* cells and visualizing the *fru P1 ∩ DIP-α* neurons. We test three isoforms of Fru^M^ (*UAS-Fru^MA^, UAS-Fru^MB^*, and *UAS-Fru^MC^*) and find they could effectively produce Fru^M^ in the expected *DIP-α* pattern (**Supplemental Table 7**). Overexpression of Fru^MB^ and Fru^MC^ has large phenotypic impacts, whereas Fru^MA^ does not, consistent with previous functional studies of the Fru^M^ isoforms (Nojima *et al*. 2014; von Philipsborn *et al*. 2014). Overexpression of Fru^MB^ results in a higher frequency of the “M” shaped projection pattern in males (curved dotted-line, “M”-like, **Figure 8E-F, subpanels II**), while the “U” shaped SEG projection is not observed as frequently (“U”-like, **Figure 8E-F, subpanels IV**). The density of the neuronal projects in the AbG is also reduced. In females, the lateral ascending neuronal projection in the VLP region is observed more frequently (**Figure 8E-F, III**). The overexpression of Fru^MC^ leads to substantial reduction of *fru P1 ∩ DIP-α* intersecting neurons in both males and females (**Figure 8G-H, subpanels III**), which could be due to a loss of neurons and/or their projects. The phenotype could also be due to reduced *DIP-α-Gal4* expression, given overexpression of Fru^M^ was previously shown to reduce expression of some IgSFs (Dalton *et al*. 2013).

A loss of the Fru^MC^ isoform, only in *fru P1* neurons, has less strong phenotypic consequences (*fru^FLP^/fru^ΔC^*; **Figure 8I-J**). In males, the SMP projections appear more female-like and there is an increase in neurons with a lateral projection, due to loss of the Fru^MC^ isoform. Therefore, overexpressing Fru^MC^ isoform in the broad *DIP-α-Gal4* pattern impacts *fru P1 ∩ DIP-α* neurons more substantially than loss of Fru^MC^ isoform in only *fru P1* expressing neurons. This suggests that the wildtype Fru^MC^ spatial expression pattern is critical for function. Furthermore, if we limit the overexpression of Tra^F^ and Fru^M^ to only *fru P1 ∩ DIP-α* neurons using an additional transgene (*tub*>*GAL80*>), we also see phenotypes that are less severe than observed when overexpression is in all *DIP-α* neurons (see **Supplemental Figure 4**). Overall, quantitative and spatial changes in the expression of sex hierarchy genes alters the sexually dimorphic *fru P1 ∩ DIP-α* patterns. This demonstrates that sex differences in morphology are downstream of sex hierarchy regulation, through both cell autonomous and non-autonomous mechanisms.

### Knockdown of *DIP-ε* in *fru P1 ∩ DIP-α* co-expressing neurons alters the expression patterns

To determine the functional roles of *dprs/DIPs* in *fru P1*-expressing neurons, we conduct an RNAi and over-expressor screen. We use the *DIP-α* and *DIP-δ* drivers, given that they have the most restricted intersecting expression patterns, which facilitates visually identifying altered patterns in *fru P1 ∩ DIP* neurons. Here, the *DIP-Gal4* drives expression of an RNAi or over-expressor transgene of other *dprs/DIPs*. It should be noted that while these *fru P1 ∩ DIP* intersecting patterns are highly restricted, the *DIP-Gal4* patterns that drive the perturbation are broader (see **Supplemental Table 4**). Out of the 36 genotypes screened, only one perturbation robustly alters the *fru P1 ∩ DIP* expression pattern (**Supplemental Table 5**). Knocking down *DIP-ε* in all *DIP-α* neurons changes the *fru P1 ∩ DIP-α* pattern (**Figure 9**). Males show a significant loss of neuronal projections that have “U” shaped arbors (see **Figure 9C, subpanel I**). Both males and females show a reduction of a set of descending neurons when compared to control flies expressing *RFP RNAi* (see **Figure 9C, subpanel II**). In addition, females show an enhancement of projections in the SMP region of the brain (see **Figure 9C, subpanel III**). These enhanced SMP projections have longer and more extensive projections that are not observed in males. Given that no other *dpr* or *DIP RNAi* perturbation shows these three phenotypes, suggests that they are specific to the *DIP-ε* perturbation (**Supplemental Table 5**). No obvious morphological changes are observed in the ventral nerve cord.

**Figure 9.**
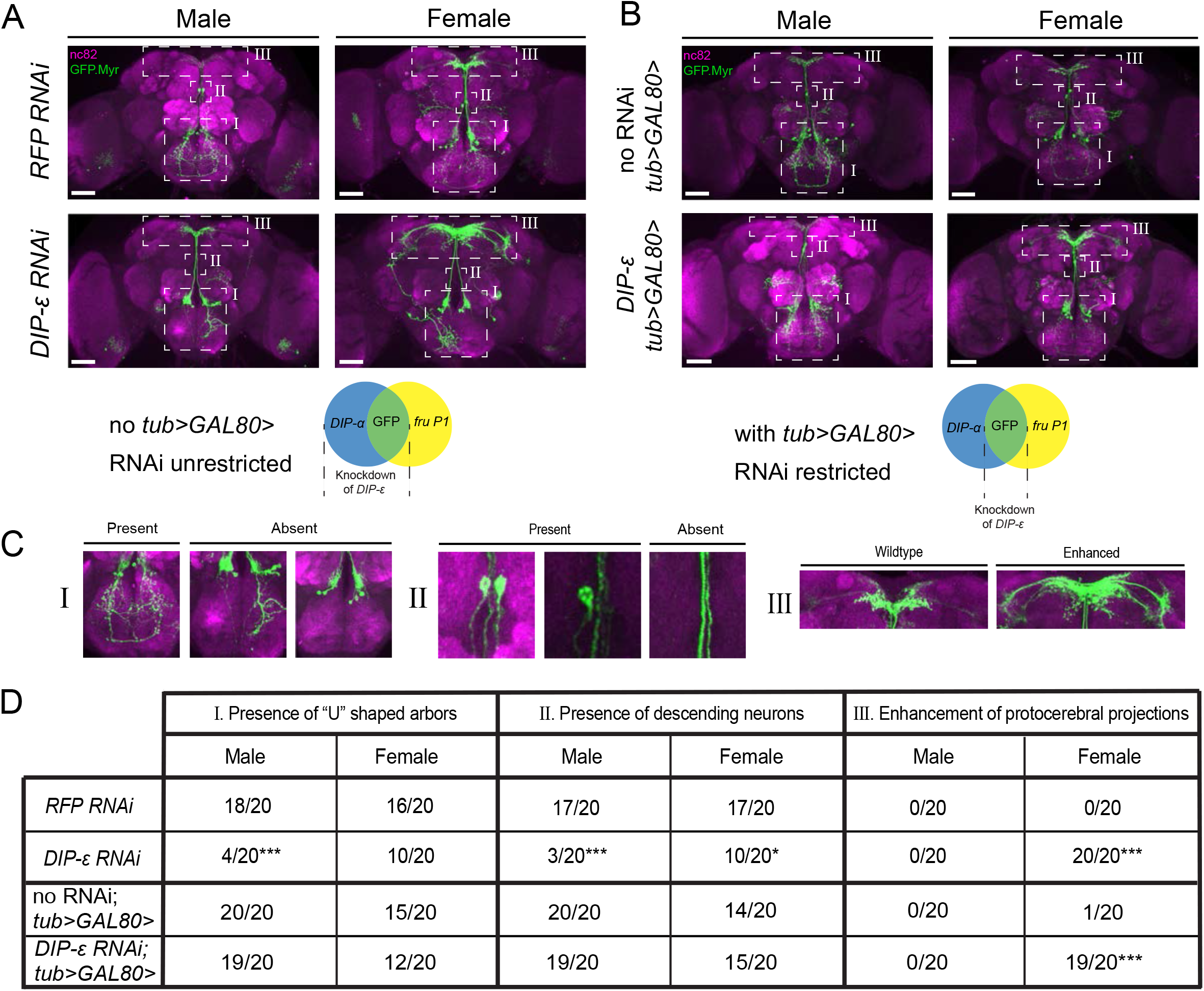
RNAi mediated knockdown of *DIP-ε* in *fru P1* ∩ *DIP-α* neurons results in perturbations. Maximum intensity projections of brains of 4-7 days old adult flies showing *fru P1* ∩ *DIP-α* neurons stained with anti-GFP (green; rabbit α-GFP Alexa Flour 488) and the neuropil marker nc82 (magenta; mouse α-nc82, Alexa Flour 633). (**A**) *fru P1* ∩ *DIP-α* neurons with *DIP-ε* or *RFP* (control) knockdown in all *DIP-α* expressing neurons. Genotypes are *DIP-α Gal4; UAS > stop > GFP.Myr /RNAi; fru^FLP^>* / + with RNAi indicating either *RFP* or *DIP-ε* RNAi. (**B**) *fru P1* ∩ *DIP-α* neurons with *DIP-ε* or no knockdown (control) restricted to only the visualized neurons (GFP+) through use of *tub>GAL80>*. Genotypes are *DIP-α Gal4; UAS > stop > GFP.Myr /RNAi; fru^FLP^ / tub>GAL80>* with RNAi indicating either *DIP-ε* or no RNAi. The neuronal populations with RNAi expression are illustrated in the Venn diagrams. White dashed boxes indicate phenotypes of interest, which are located in (**C**) and include (**subpanel I**) presence of the U-shaped arbors, (**subpanel II**) presence of at least one descending neuron, and (**subpanel III**) enhancement of protocerebral projections. All phenotypes were scored blind and are quantified in (**D**). Statistical significance in between control flies and *DIP-ε* RNAi flies was evaluated by the Fisher’s exact test. In this figure, signicance is indicated as follows: *P < 0.05, **P < 0.01, ***P < 0.001. n=20 brains for each category. Magnification is 20x and scale bars represent 50 μM.

We next examine the phenotypes when the *DIP-ε RNAi* knockdown is limited to only the *fru P1 ∩ DIP-α* co-expressing neurons, rather than all *DIP-α* neurons. We continue to use the genetic intersecting approach to visualize the neurons with GFP. To restrict expression of *DIP-ε RNAi* to *fru P1 ∩ DIP-α* neurons we use an additional construct (*tub>GAL80>*), such that Gal4 is now only transcriptionally active in *fru P1 ∩ DIP-α* (**Figure 9B**). Males no longer show a significant reduction of the “U” shaped projections, and neither sex shows a significant reduction of descending neurons (**Figure 9**). This suggests that these phenotypes are due to reduction of *DIP-ε* outside of *fru P1 ∩ DIP-α* neurons, in a non-cell-autonomous manner. Conversely, females still have the enhanced projections in the protocerebrum. This suggests that this phenotype is cell autonomous and driven by a reduction in *DIP-ε* expression inside the *fru P1* **∩** *DIP-α* neurons. The results are consistent with the observation that both *fru P1* ∩ *DIP-α* and *DIP-ε* are expressed in similar patterns in the SMP (**Figure 3**) and so it is not unexpected that expression of *DIP-ε RNAi* can have a functional impact in *fru P1* ∩ *DIP-α*. Taken together, these results demonstrate that *DIP-ε* plays a critical role in establishing wildtype *fru P1* neuronal patterns, in both a cell-autonomous and non-cell-autonomous manner.

### Single cell mRNA sequencing analysis in male *fru P1*-expressing cells

To examine the repertoires of *dprs/DIPs* expressed in individual *fru P1* neurons, we perform single cell sequencing (10X Genomics). The analysis is performed on male central nervous system tissues (48-hour pupal stage), from flies that expressed membrane-bound GFP in *fru P1* neurons. We chose this developmental stage to gain further insight into how the *dprs/DIPs* direct development of *fru P1* neurons, as this is the stage where Fru^M^ has peak expression (in ~2,000 neurons, Lee *et al*. 2000). The matrix of the single cell sequencing data is filtered to identify the *fru P1* neurons, based on detection of the membrane-bound GFP mRNA, which resulted in 5,621 cells for analysis. We perform normalization and data scaling using all genes in the matrix, for data from the *fru P1* neurons. We find that all *fru P1* neurons express at least one *dpr/DIP*. Then a principle component analysis (PCA) is performed using only *dpr*/*DIP* gene expression, and the dimensionality is reduced with the UMAP algorithm (McInnes and Healy, 2018 arXiv:1802.03426 and Stuart *et al*. 2019). Cells with similar *dpr/DIP* expression will cluster closely with one another in the UMAP plot (**Figure 10A**). A visual inspection of the UMAP plot reveals that the patterns of *dpr/DIP* expression are not distinct enough to generate highly refined clusters that have large separation in the UMAP plot.

**Figure 10.**
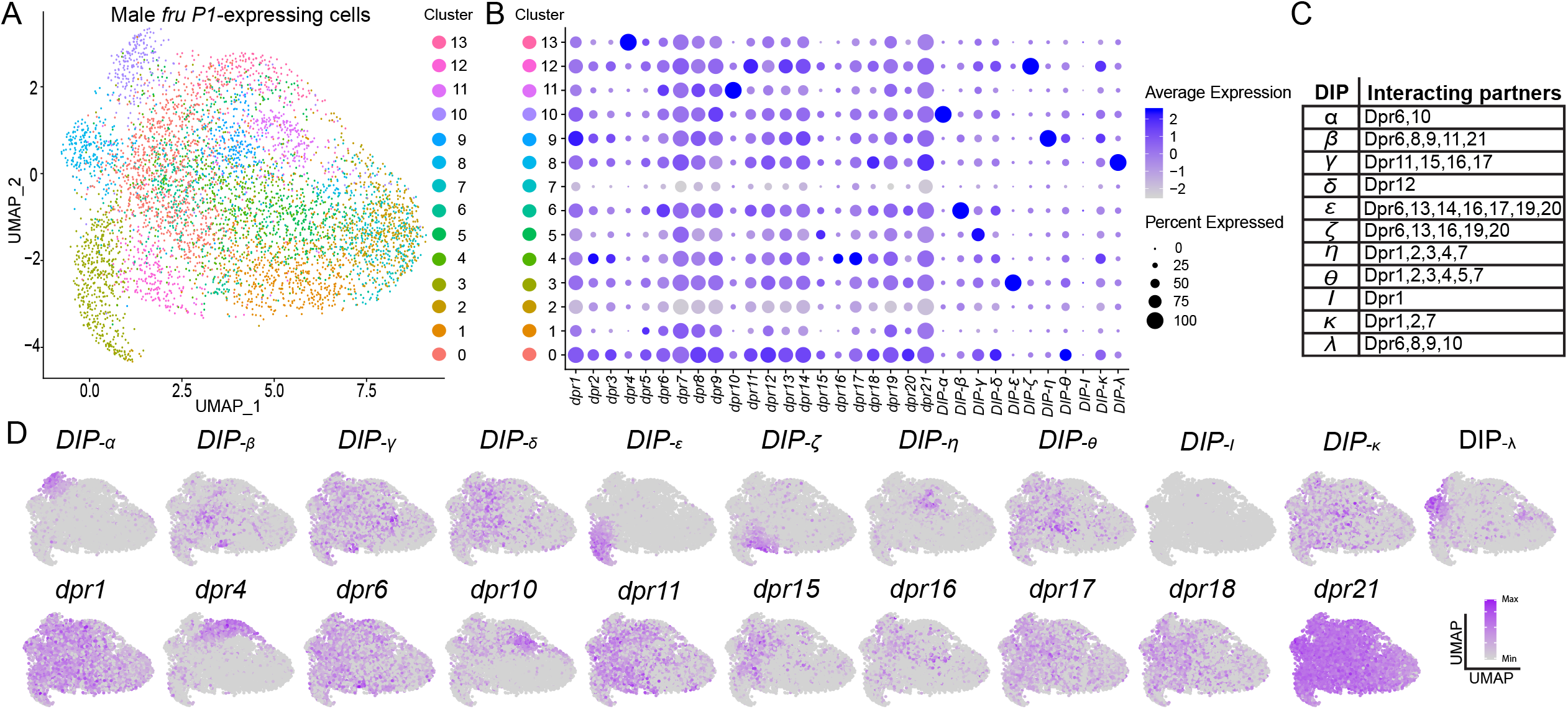
Single cell RNA-sequencing analysis of *dpr/DIP* expression analysis in male *fru P1*-expressing cells. **(A)** UMAP plot of *5,621 fru P1*-expressing nervous system cells, isolated from male tissue 48hr after puparium formation. The data are clustered on *dpr/DIP* gene expression. **(B)** Dot plot showing the expression of *dpr/DIP* genes across all clusters identified in UMAP. Dot diameter indicates the fraction of cells expressing each gene in each cluster, as shown in legend. Color intensity indicates the average normalized expression levels. **(C)** Heterophilic interactions between DIPs and Dprs. The Dpr-DIP interactions are previously described (Carrillo *et al*. 2015). The interacting partners for DIP-κ and DIP-λ are previously described (Cosmanescu *et al*. 2018), as they were not part of the Carrillo 2015 study. **(D)** A subset of expression visualization of *DIPs* (top row) and subset of *dprs* (bottom row) in the UMAP-clustered cells. *dpr* or *DIP*-positive cells are labeled purple and color intensity is proportional to log normalized expression level shown in legend. The UMAP for all *dprs/DIPs* is provided (**Supplemental Figure 5**). The numerical expression values are in **Supplemental Table 6**.

For each cluster, we next determine if a combination of *dprs/DIPs* are largely responsible for each cluster identity. We examine the average expression and the percent of cells with expression of each *dpr/DIP* in each cluster (**Figure 10B**). We find that the majority of the *DIPs* have high average expression in one cluster, with a large percent of the cells in the cluster having expression. This is distinct from the majority of the *dprs*, where the average expression and the percent of cells that express the *dpr* is more moderate and similar across many clusters. It does not appear that most of the clusters are due to co-expression of Dpr/DIP interacting partners (**Figure 10C**), based on a visual inspection. Furthermore, the distribution of the expression patterns overlaid on the UMAP plot for each *dpr/DIP* also shows that the *DIP*s have more restricted expression. For example, *dpr21* is broadly detected across the UMAP plot, whereas *DIP-α* has restricted expression in cluster 10, at the upper left-hand side of the UMAP plot (subset of expression patterns in **Figure 10D;** for all *dprs/DIPs* see **Supplemental Figure 5**). This suggests that *in vivo*, DIPs may have different functional roles, compared to Dprs, in terms of directing synaptic specificity or cell adhesion properties of the neuron.

We also generate a dendrogram, by hierarchical clustering, to visualize which *dprs* and *DIPs* have the most similar expression patterns, using the same normalized and scaled gene expression data matrix that is used to generate the UMAP plot (**Supplemental Table 6**). We find that some *dprs* and *DIPs* that have shared interacting partners have the most similar expression to each other. This includes the following pairs: *dpr2* and *dpr3; dpr6* and *dpr10; dpr16* and *dpr17;* and *DIP-ζ* and *DIP-ε*. We find that *DIP-α, DIP-ι* and *cDIP* have the most distinct expression patterns from the rest of the interactome, which may be due to low number of cells in which they are detected (see Upset plot described below; **Supplemental Table 6**). For some neurons, co-expression of *dprs* and *DIPs* with the same interacting partners may be a mechanism to generate different adhesion properties.

Next, we examine the number of different combinations of *dpr* and *DIP* expression repertoires. To do this analysis a gene is considered expressed within a neuron if the normalized and scaled expression data value is >1, thus excluding those with stochastic expression detection due to low expression levels (403 neurons do not have *dpr/DIP* expression based on this criterion; 5,218 neurons remain). There are 458 neurons that express only one *dpr* or *DIP*. The range of neurons that express 2-8 *dprs* and *DIPs* is between 451-653 neurons (4,024 total); that express 9-11 *dprs* and *DIPs* is between 105-332 neurons (657 total); and that express 12-15 *dprs* and *DIPs* is between 5-45 neurons (79 total; **Supplemental Table 6**). Next, we look at the number of neurons with the same expression repertoire. This can be ascertained using an “Upset” plot, which is conceptually similar to a Venn Diagram, but accommodates a large number of conditions, which here are the 5,218 single neuron expression repertoires. The majority of expression repertoires that are detected in more than one neuron are those for which the neuron only expresses one *dpr* or *DIP* (single dots on bottom of Upset Plot, 457 neurons; **Supplemental Table 6**). There were also 466 neurons that had shared co-expression combinations due to expression of 2-5 *dprs* and *DIPs*. The majority of *fru P1* neurons had a unique repertoire of *dpr/DIP* expression (4,295 neurons), due to expression of different combinations of 2-15 *dprs* and *DIPs* (**Supplemental Table 6)**. In the developing *fru P1* neurons, this singular and co-expression of *dprs* and *DIPs* provides a mechanism to generate different connectivity properties for each cell.

## Discussion

Based on our systematic reevaluation of previous genomic data sets and the microscopy results presented, we show that *dprs/DIPs* are regulated by Fru^M^ and expressed in *fru P1* neurons in both males and females (**Figures 1–3**). The expression pattern for each *fru P1* ∩ *dpr/DIP* genotype is unique, though many genotypes have expression in the same brain regions, including the lateral protocerebral complex, mushroom body, antennal lobe, tritocerebral loop, mesothoracic triangle and abdominal ganglion (**Figure 7**), which are regions that were previously shown to be among those with the most pronounced sexual dimorphism in *fru P1* neurons (Cachero *et al*. 2010; Yu *et al*. 2010). Furthermore, while the patterns for each genotype are similar between males and females, we find sexual dimorphism in some of the projection patterns and in neuron numbers (**Figures 2,3 and 7**). Given that the *dprs/DIPs* are not sex-specifically expressed, this suggests that their role in generating sexual dimorphism may be quantitative, due to sexual dimorphism in expression levels or differences in the number of neurons in which they are expressed in a given region.

We find that activating and silencing the subsets of neurons defined by each *fru P1* ∩ *dpr/DIP* genotype differentially impacts male courtship behaviors, with the results highlighting that the activity of different combinations of neurons can generate a similar behavioral outcome. This analysis provides further insight into how similar behavioral outcomes can be generated in different ethological contexts, through the integration of information across many different neuronal subtypes. An examination of the similarities of *fru P1* ∩ *dpr/DIP* expression patterns and behavioral outcomes suggests that interactions between the mushroom body and lateral protocerebral complex are critical to reach a certain threshold of activation for male courtship behaviors, in both the male-female and male-alone paradigm, given that the genotypes with expression in those two regions had the most consistent and robust behavioral phenotypes. Interactions between neurons in these two regions have previously been proposed to integrate disparate sensory information and behavioral experiences, to direct courtship outcomes (Yu *et al*. 2010).

A higher resolution analysis of *fru P1 ∩ DIP-α* neurons found additional sexual dimorphism in projections and neuron number that are downstream of the sex hierarchy. Regulation by the sex hierarchy of *fru P1 ∩ DIP-α* neurons is both cell-autonomous and cell non-autonomous. These result point to the importance of understanding the development and function of *fru P1* neurons in a broad context, taking into account interactions with both *fru P1* and non-*fru P1* neurons. Furthermore, an RNAi and overexpression screen show there is functional redundancy in patterning, with only *DIP-ε RNAi* generating phenotypes. A single cell RNA-seq analysis of *fru P1* neurons shows that *dprs/DIPs* are expressed in every *fru P1* neuron, with the majority having a unique expression combination. The UMAP cluster analysis shows that generally *DIPs* have high average expression in a small set of neurons, whereas *dprs* have more moderate expression across a larger set of neurons, suggesting that they may have different functional roles.

### Role of Dprs and DIPs in sexual dimorphism of *fru P1* neurons

In the optic lobe, antennal lobe and neuromuscular junction, genetic analyses have demonstrated that Dprs/DIPs have a role in synaptic specificity and connectivity, with Dpr-DIP interactome partner pairs mediating these critical functions (Carrillo *et al*. 2015; Tan *et al*. 2015; Barish *et al*. 2018; Xu *et al*. 2018; Ashley *et al*. 2019; Courgeon and Desplan 2019; Menon *et al*. 2019; Venkatasubramanian *et al*. 2019; Xu *et al*. 2019). Our screen to identify morphological or synaptic changes in *fru P1 ∩ DIP-α* and *DIP-δ* neurons, using *dpr/DIP* RNAi or overexpression transgenes identified only one perturbation with an impact; reduction of *DIP-ε* by RNAi on the *fru P1 ∩ DIP-α* pattern, with both cell autonomous and non-autonomous roles. This suggests that there is sufficient redundancy that removal or addition of one member of the Dpr/DIP interactome cannot change patterning robustly. It is possible that this is due to overall patterning by the other Dpr/DIP interactome pairs, given how many different combinations of *dpr/DIP* genes are expressed in the majority of *fru P1* neurons, based on the single cell sequencing data. DIP-ε interacts with a large number of Dprs, which may be one of the reasons a reduction of DIP-ε results in morphological changes. We found that some Dprs that interact with the same DIP are expressed in the same brain regions (**Figure 7**), and/or are detected in the same neurons (**Supplemental Table 6**), consistent with the idea of redundancy. This could also be due to other members of the IgSF that were identified by our genomic-scale screens as expressed in *fru P1* neurons, or other guidance molecules. The enhanced set of projections in the superior medial protocerebrum region of the brain due to reduced *DIP-ε* is reminiscent of the synaptic targeting phenotypes seen in the optic lobe due to *dpr/DIP* perturbations (Carrillo *et al*. 2015; Tan *et al*. 2015; Courgeon and Desplan 2019; Menon *et al*. 2019; Xu *et al*. 2019), which supports a role of *dprs/DIPs* in the development of *fru P1* neuroanatomical projection patterns and/or synaptic targets.

Future studies that are performed with genetic tools that yield more penetrant phenotypes than RNAi, including Crispr/Cas9 generated alleles, will likely reveal additional roles for Dprs/DIPs. Furthermore, Crispr/Cas9 gene knock-out approaches can target multiple *dprs/DIPs*, allowing one to test for functional redundancy. Additional analyses to determine the subcellular localization of each Dpr/DIP will also be important to understand their roles in the nervous system, especially to determine if they are present in synaptic termini and dendrites, which would be consistent with a role of synaptic specificity. It is clear that higher resolution analyses of the *fru P1 ∩ DIP-α* pattern reveal more sexual dimorphism, so additional analysis of other genotypes at this resolution will be important, including determining developmental patterns to gain insight into mechanisms that underlie sexual dimorphism. Furthermore, our expression data reveal expression beyond development, well into adult stages. Adult roles of the *dprs/DIPs* may include mediating neuronal connectivity changes due to reproductive experiences. The results of the single-cell RNA sequencing analyses show that the majority of expression repertoires of the *dpr* and *DIP* genes is distinct within each *fru P1* neuron. Additionally, we find that most *DIPs* have high expression in a small set of neurons, and most *dprs* have moderate expression across a larger set of neurons. One possibility is that DIP expression in a neuron provides more information about cell fate identity, because it is more restricted and at higher levels.

### *fru P1* ∩ *dpr/DIP* neurons and male courtship behaviors

In this study each *fru P1* ∩ *dpr/DIP* genotype has different expression patterns across the nervous system, allowing us to ascertain if different combinations of neurons are critical for a behavioral outcome. We found that genotypes that had neuronal activation in both mushroom bodies and the lateral protocerebral complex had the most consistent observation of atypical behaviors and overall courtship in both the male-female and male-alone courtship behavioral studies. While there has been an impressive effort to map functions onto small subsets of neurons (Robie *et al*. 2017), our results suggest that it will also be important to understand the roles of different combinations of neurons to fully understand behavioral outcomes. This will facilitate understanding of how different sensory and courtship experiences impart physiological changes to direct behavior. Furthermore, these activation experiments may also reveal insights about evolution of behavior. In some Drosophila species, males perform a double wing extension during courtship (reviewed in Anholt *et al*. 2020). We observe double wing extension in several genotypes in the neuronal activation experiments, suggesting that changing levels of neuronal activity are a way to evolve a new behavior. While this study focused on male reproductive behaviors, it will also be interesting to examine the role of *fru P1* ∩ *dpr/DIP* neurons on female behavioral outcomes.

## Conclusions

Over the last several years genomic studies have pointed to a role of the *dprs/DIPs* in *fru P1* neurons (Goldman and Arbeitman 2007; Dalton *et al*. 2013; Neville *et al*. 2014; Vernes 2014; Newell *et al*. 2016). Indeed, our early study showed that *dpr1* had a role in courtship gating, or the timing of the steps that the male performs (Goldman and Arbeitman 2007). Until recently, a systematic analysis of the role of *dprs/DIPs* in *fru P1* neurons was not possible. Future studies aimed at a systematic analysis of the Dpr/DIP interactome will further elucidate the role of these cell adhesion molecules in terms of specifying neuroanatomy and also as powerful tool to gain insight into the functions of different sets of *fru P1* neurons.

## Materials and methods

### Fly husbandry and stocks

All flies were raised at 25°C on a 12:12 hours light-dark cycle. The flies were grown using standard cornmeal food media (33 L H2O, 237 g Agar, 825 g dried deactivated yeast, 1560 g cornmeal, 3300 g dextrose, 52.5 g Tegosept in 270 ml 95% ethanol and 60 ml Propionic acid). A list of Drosophila strains is provided (**Supplemental Table 7**).

### Immunohistochemistry and confocal microscopy

Brain and ventral nerve cord (VNC) tissues were dissected from animals that were either 0-24 hour adults, or 4-7-day adults. Samples were dissected in 1x Phosphate Buffered Saline (PBS; 140 mM NaCl, 10 mM phosphate buffer, and 3 mM KCl, pH 7.4) and immediately transferred to fix (4% paraformaldehyde, 1x PBS) for 25 minutes at room temperature. Samples were washed for 5 minutes with 1x PBS, three times. The tissue was then permeabilized with TNT (0.1 M Tris-HCl [pH 7.4], 0.3 M NaCl, 0.5% Triton X-100), for 15 minutes, followed by two additional 5 minute TNT washes. The tissue was rinsed in 1x PBS, and then Image-iT^™^ FX Signal Enhancer (Invitrogen) was applied for 25 minutes. Finally, the tissue was washed in TNT for two washes of 5 minutes each. Diluted primary antibody in TNT was applied, and samples were incubated overnight at 4°C. Next, the tissue was washed six times in TNT for 5 minutes each, and then secondary antibody diluted in TNT and applied. The samples were then incubated for 2 hours at room temperature or overnight at 4°C. Following this incubation, samples were washed six times in TNT for 5 minutes each and then mounted in Secureseal^™^ Image Spacers (Electron Microscopy Services), on glass slides with VectaShield^®^ Mounting Medium (Vector Laboratories; H-1000), and covered with #1.5 coverslips. Primary antibodies were used in the following dilutions, as indicated in the figure legends: mouse α-nc82 (1:20; Developmental Studies Hybridoma Bank, AB_2314866), rabbit α-Myc (1:6050; abcam, ab9106), rabbit α-GFP Alexa Fluor 488 (1:600; Invitrogen, A21311). Secondary antibodies were used in the following dilutions: goat α-rabbit Alexa Fluor 568 (1:500; Invitrogen, A11036), goat α-mouse Alexa Fluor 633 (1:500; Invitrogen A21052). For labeling of three MCFO markers (FLAG, V5, and HA), brains and VNCs samples were dissected and stained by following the method modified from Nern et al (Nern *et al*. 2015). The primary antibodies rabbit α-HA (1:300; Cell Signaling, 3724S), mouse α-FLAG (1:500; Sigma, F1804), and rabbit α-V5 DyLight 549 (Rockland, 600-442-378), and the secondary antibodies goat α-rabbit Alexa Fluor 633 (1:500; Invitrogen A21071) and goat α-mouse Alexa Fluor 488 (1:500; Invitrogen A11001) were used. All the antibodies were diluted in TNT.

Images were acquired on a Zeiss LSM 700 confocal microscope with a 20x objective and bidirectional scanning. The interval of each slice was set as 1.0 μm. Zeiss Zen software (Black edition, 2012) was used to make adjustments to laser power and detector gain to enhance the signal to noise ratio.

### Live tissue staining

Conditioned media containing the extracellular domain (ECD) of the Dpr/DIPs was generated by transfecting *Drosophila* S2 cells with DNA plasmids, as previously described (Ozkan *et al*. 2013). S2 cells were seeded at 2×10^6^ per 6cm plate in 4 mL S2 medium (Lonza). One hour after plating, S2 cells were transfected with 1 ug plasmid DNA using the Effectene reagent kit (Qiagen). The plasmid DNA which contains cDNA of FLAG-tagged ECDs are under the metallothionein promoter control. Therefore, 1 mM CuSO4 was used to induce ECD expression 18-hours after plasmid DNA transfection. Conditioned media were collected after 3days of 1mM CuSO4 induction. S2 cells were removed by 10 minutes of gentle spinning at 1,500g and the supernatant was further spun through an Amicon Ultra-4 Centrifugal Filter, with 100 kDa cut-off, to concentrate the conditioned media containing the ECD. The supernatant was stored at 4°C with 0.02% sodium azide and protease inhibitors (Sigma, P8849).

For live tissue staining, *Drosophila* central nervous systems tissue were dissected in S2 medium and then incubated with conditioned S2 medium for 18 hours at 4°C on a rotating platform. After the incubation, tissues were washed with S2 medium and fixed with 4% paraformaldehyde in 1x PBS for 45 minutes. After fixation, tissues were further washed with two times of 1x PBS and two times with TNT. ECD binding was detected through overnight incubation of 1:500 of anti-FLAG antibody (Sigma) at 4°C. Goat anti-mouse Alexa Flour 488 (1:500) was used as the secondary antibody. Three times of TNT wash were performed before slide and imaging, as described above.

### Image analysis and quantification of *fru P1* ∩ Dpr/DIP neurons

Brain and VNC confocal images of 4-7-day old male or female adults were analyzed for the presence of certain morphological features and cell body numbers of select neurons. The images were scored blind, in randomized batches, by three independent people. The analysis was performed using Fiji-ImageJ 14.1, with the cell counter Janelia version 1.47h plugin. To determine which regions to analyze, the following criteria were used: 1) regions that had sexually dimorphic structures, 2) were present in many of the different genotypes, and/or 3) are known to be important for reproductive behaviors. A template of example images, with regions indicated, was used to ensure accurate and similar image analyses across all researchers (**Supplemental Table 2**). As a test to ensure accuracy of scoring across the three individuals, a round-robin scoring design was employed, with each image scored by three individuals, for a subset of 26 images, which showed high concordance. The raw cell count numbers and morphological observations were recorded in excel, compiled and then unblinded (**Supplemental Table 2**).

### Generation of heatmaps

Heatmaps and correlation plots of the image analysis data were generated using Morpheus (Broad Institute; https://software.broadinstitute.org/morpheus). For features that were scored as present or absent, a value of 0 or 1 was calculated as the number of samples with the feature present divided by total number of samples. For the cell count data, the replicate data was averaged, and then all data was divided by the highest value for that cell count feature, so all data were between 0 and 1. The hierarchal cluster heatmap was made using the following parameters: one minus spearman rank correlation as the metric, average for linkage method, and clustering by the columns (data for each *dpr/DIP*). The correlation heatmaps were created using the Morpheus similarity matrix tools, using the following parameters: spearman rank as the metric, computed for the columns.

### Courtship behavior assays and analyses

For all behavior, male flies were collected 0-6 hours post-eclosion, housed individually in small vials, and aged for 4-7 days. Canton S virgin females (*white*) were also collected 0-6 hours post-eclosion, and aged for 4-7 days in groups to be used as female targets for courtship with males containing the *UAS > stop > TrpA1:myc* transgene. Canton S virgin females were collected and kept in a similar manner to be used for courtship with male flies containing the *UAS > stop > TNTE/TNTQ* transgenes. Flies were kept in a 25°C incubator on a 12:12 hour light:dark cycle, unless otherwise noted. Courtship chambers were placed on a temperature-controlled metal block at 25°C and videos were recorded between ZT 5-10, in a 10-mm chamber for ten minutes, or until successful copulation occurred, whichever came first. For courtship using male flies harboring the *UAS > stop > TrpA1:myc* transgene, the male flies were reared and housed in a 19°C incubator, on a 12:12 light:dark cycle, so the Trp channel would not be activated. Courtship chambers were placed on a temperature-controlled metal block for ten minutes prior to the courtship assay, at either 20°C or 32°C.

The courtship video recordings were analyzed using The Observer^®^ XT (Noldus) (version 14.0), with an n=14-16 for male-female behavior and n=10 for male alone behavior. Coded behaviors included: following (a start-stop event defined as any time the male is oriented towards the female and is less than half a chamber distance away from the female), wing extension (a start-stop event defined as any time one wing is extended from the fly and is vibrating), double wing extension (a start-stop event when both wings are extended from the body and are vibrating), abdominal bending (a start-stop event when the abdomen is curled under and is not thrusting or is not in the correct position to copulate with the female), motor defect (a start-stop event when the male falls onto his back and is unable to right himself), attempted copulation (a point event when the male attempts to copulate with the female but is not successful), and successful copulation (a point event when the male is able to attach and successfully copulate with the female).

These data were graphed and analyzed using the JMP^®^ Pro 14.0.0 statistical software. A non-parametric Wilcoxon test was used to compare differences between the control and experimental temperature (for TrpA1 experiments) or between control and experimental strains (for TNT experiments), for the data for which an index is calculated. The unpaired *t-test* was used to determine significant differences between experimental and control conditions, with the same *dpr/DIP-Gal4*, to determine if the number of attempted copulations were different (test assumes equal variance).

### *Drosophila* activity monitor behavioral assay

Males were collected 0-6 hours post-eclosion and aged for three days in a 25°C incubator on a 12:12 hour light:dark cycle. On day three, they were individually loaded into 5 × 65mm glass tubes (Trikinetics Inc.), plugged on one end with standard cornmeal food media dipped in paraffin wax to seal. The non-food end was sealed with parafilm, with small air holes. The vials were loaded into *Drosophila* activity monitors (TriKinetics Inc.), and placed in a 25°C incubator in 12:12 hour light:dark. Each condition was run for five days. The data from the first day of activity was not used in the analysis, as flies were recovering from CO2 anesthesia. Activity was measured as the number of beam breaks and collected in five-minute bins. Beam crossings were summed over the 24-hour period from day 5 ZT0 (lights-on) to day 6 ZT0 per individual fly (**Supplemental Table 3**). These data were graphed and analyzed using the JMP^®^ Pro 14.0.0 statistical software (**Supplemental Figure 6**). A non-parametric Wilcoxon test was used to compare differences between the control (TNTQ) and experimental (TNTE) strains with the same *dpr/DIP-Gal4*.

### Image analyses of RNAi and over-expression perturbations

#### Functional roles of *dpr/DIPs*

Initially, several different combinations of one *dpr/DIP-Gal4* driver, and either a UAS-RNAi *dpr/DIP*, or a *UAS-dpr/DIP* expression transgene were assayed, using the intersectional genetic approach for visualization of small sets of *fru P1*-expressing neurons (**Figure 1C; Supplemental Table 5**). For the RNAi screen, parents laid eggs at 25°C for 2-3 days, and then the vials with eggs were transferred to 29°C, to increase effectiveness of RNAi constructs. For the over-expression screen, flies were raised at 25°C. Staining and confocal imaging was performed as described above. Through this initial screen, we found that knocking down *DIP-ε* in *DIP-α* **∩** *fru P1* neurons at 4-7 days was the only condition to yield a robust phenotype. Knockdowns were analyzed, with *DIP-ε* or *RFP* RNAi active in all *DIP-α* expressing cells. In addition, knockdowns were restricted to the visualized *fru P1* **∩** *DIP-α* neurons with the use of *tub>GAL80>*.

The *fru P1* **∩** *DIP-α* neurons were analyzed blind in 20 brains, in male and female controls and mutants, to determine the effect of *DIP-ε* knockdown on neuronal morphology (**Supplemental Table 5**). The presence or absence of morphological features were compared within sex between *DIP-ε* knockdowns and the corresponding control using a Fisher’s exact test (R version 3.5.1, R Core Team, 2019).

#### Sex hierarchy perturbations

The *DIP-α* subset of *fru P1*-expressing neurons were further analyzed to determine the impact of sex hierarchy perturbations. Flies bearing RNAi and over-expressor constructs were raised to 4-7-day old adults, stained, and imaged as described above. Both RNAi knockdown and overexpression experiments were first performed in all *DIP-α* expressing cells. In addition, the over-expressors were also restricted to the visualized *fru P1* **∩** *DIP-α* neurons with the use of *tub>GAL80>*.

The *fru P1* **∩** *DIP-α* neuronal patterns were analyzed blind in at least 15 brains and ventral nerve cords, in males and females, to determine the effect of sex hierarchy perturbations on neuronal morphology (**Supplemental Table 4**), for a set of morphological features (**Supplemental Table 4**). The ratios of different types of the morphological features and presence or absence of morphological features were compared within sex, between sex hierarchy perturbation groups and the corresponding controls using Fisher’s exact test (tests were conducted in R version 3.5.1, R Core Team, 2019).

### Dissociation of CNS for single cell mRNA sequencing analyses

Twenty freshly dissected male brains and ventral nerve cords, from 48 hour after puparium formation (APF) stage, were used. The flies had expression of membrane-bound GFP in *fru P1*-expressing neurons and were the following genotype: w[*]; P{y[+t7.7] w[+mC]=10XUAS-IVS-mCD8::GFP}attP40*/UAS-Gal4,ofru P1-Gal4/+*. The tissue was dissected in cold Schneider 2 Drosophila culture medium (S2 medium, Gibco) and transferred to a LoBind tube (Eppendorf), containing 200μl of S2 medium. The tissue was centrifuged at 500g for 5 minutes, and then was washed with 300μl of EBSS (Earle’s Balanced Salt Solution), and centrifuged again at 500g for 5 minutes. After centrifugation, the supernatant was replaced with 100μL of papain for disassociation (50 units/ml, Worthington) diluted in EBSS. Brains were dissociated at 25°C in a LoBind tube for 30 min., with pipette mixing to reinforce dissociation every 3 min with a P200 tip during the first 15 minutes, and a P10 tip for the final 15 min. Cells were washed twice with 700μl cold S2 medium containing 10% FBS (Gibco) and centrifuged at 700g for 10 minutes to quench the papain. Cell suspensions were passed through a 30μM pre-separation filter (MiltenyiBiotech). Cell viability and concentration were assessed by hemocytometer using Trypan blue.

### 10xGenomics library preparation and sequencing

Single-cell libraries were generated using Single Cell 3’ Library & Gel Bead Kit v2, Chip Kit, and the GemCode 10X Chromium instrument (10X Genomics, CA), according to the manufacturer’s protocol (Zheng *et al*. 2017). In brief, single cells were suspended in S2 medium with 10% FBS and the maximum volume of cells, 34μl, was added to a single chip channel. After the generation of nanoliter-scale <u>G</u>el bead-in-<u>EM</u>ulsions (GEMs), the mRNA in GEMs underwent reverse transcription. Next, GEMs were broken, and the single-stranded cDNA was isolated, cleaned with Cleanup Mix containing DynaBeads MyOne Silane beads (Thermo Fisher Scientific). cDNA was then amplified with the following PCR machine settings: 98°C for 3 min, 9 cycles of (98°C for 15s, 67°C for 20s), 72°C for 1 min, held at 4°C. Subsequently, the amplified cDNA was cleaned up with SPRIslect Reagent kit (Beckman Coulter), fragmented, end-repaired, A-tailed, adaptor ligated, and cleaned with SPRIselect magnetic beads between steps. This product was PCR amplified with the following PCR machine settings: 98°C for 45s, 12 cycles of (98°C for 20s, 54°C for 30s, 72°C for 20s), 72°C for 1 min, and hold at 4°C. The library was cleaned and size-selected with SPRIselect beads, followed by Pippin size selection for a 350-450bp library size range. Single cell libraries were sequenced on the Illumina NovaSeq with 150bp paired-end reads on an S2 flowcell. This produced 1,870,220,065 reads.

### Single cell data pre-processing and analysis

Raw reads were processed using the CellRanger software pipeline (v.2.1.1) “cellranger count” command to align reads to the *Drosophila melanogaster* (BDGP6.92) STAR reference genome, customized to contain the sequence for the *mCD8-GFP* cDNA. The “force-cells” command was used to call 25,000 single cells, based on the inflection point of the CellRanger barcode rank plot, a criterion for dividing single cells from empty GEM droplets (**Supplemental Table 6**). The recovered 25,000 single cells had a mean sequencing depth of 74,808 reads per cell. We detected a median of 2,118 genes per cell. The obtained feature-barcode matrix was further processed and analyzed in the R package Seurat (v3.0) (Stuart *et al*. 2019). To filter the expression matrix for high quality cells we removed cells with >5% mitochondrial transcripts (dying cells), <200 genes (empty droplets), and/or expressing more than 6000 genes (potential doublets or triplets). This filtering produced a matrix of 24,902 high quality cells which were computationally subset to the population of *fru P1*-expressing cells, based on *mCD8-GFP* expression, obtaining 5,621 cells. We next followed the Seurat “Guided clustering tutorial” for default normalization and scaling steps (https://satijalab.org/seurat/v3.0/pbmc3k_tutorial.html). Expression was normalized using the “NormalizeData” function where gene counts within each cell are divided by the total gene counts for that cell, multiplied by a scaling factor of 10000, and natural-log transformed (log1p). A linear transformation was applied to the normalized gene counts, to make genes more comparable to one another, using the default “ScaleData” function to center the mean expression to 0 and set the variance at 1. We performed a Principle Component Analysis (PCA) using only the data from 33 *dpr/DIP* genes. We used the top 20 principle components based on visual inspection of DimHeatmap outputs and the ElbowPlot. Selecting more than 20 PCs did not dramatically change our results. We then continued to follow Seurat’s standard workflow to reduce dimensionality and cluster cells using the default “FindNeighbors”, “FindClusters”, and “runUMAP” functions (resolution = 1.3).

To evaluate expression combinations of the *dpr/DIPs* within our single cells we used an UpSet plot analysis (Conway *et al*. 2017). To do this, we transposed our matrix which contained normalized, log-transformed, and scaled expression data (**Supplemental Table 6**) for *dpr*/*DIP*s for each single cell barcode and binarized the data (any expression of a *dpr* or *DIP* > 1= 1, and >1 is considered as no expression = 0, **Supplemental Table 6**). All plots generated are ordered by the highest frequency of an expression combination occurring within single cells (order.by = “freq”). A single cell expression hierarchical clustering dendrogram was produced using the normalized, log-transformed, and scaled expression data (**Supplemental Table 6**). A Pearson correlation distance measure was calculated using the factoextra (v. 1.0.7) “get_dist” function and hierarchical cluster analysis was performed using the “hclust” core R statistics function with the argument method= “average”.

## Acknowledgements

The work presented was supported by NIH grants awarded to MNA: R01GM073039, R01GM116998, R03NS090184. This work was also supported by funds from the Biomedical Sciences Department, College of Medicine, Florida State University. We are grateful for the support. We appreciate that colleagues sent Drosophila stocks (Supplemental Table 7). Stocks were also obtained from the Bloomington Drosophila Stock Center (NIH P40OD018537). Several antibodies used in this study were obtained from the Developmental Studies Hybridoma Bank, created by the NICHD of the NIH and maintained at The University of Iowa, Department of Biology, Iowa City, IA 52242. We appreciate experimental assistance from Catherina Artikis.

---

**Supplemental Figure 1.**
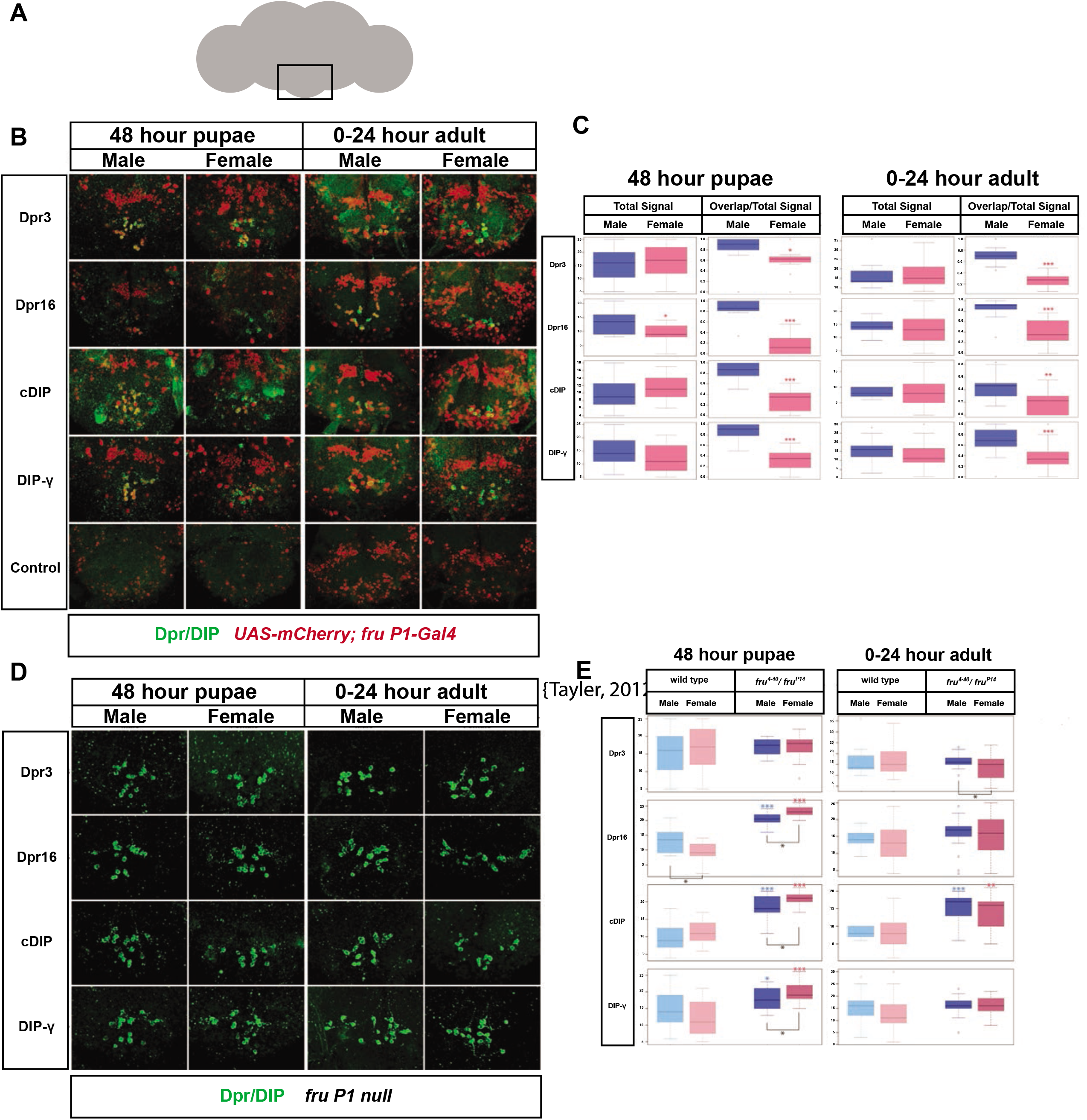
Live *in vivo* staining of Drosophila brain tissues. Staining was performed using S2 culture media from cells expressing the extracellular region of a Dpr or DIP that is tagged with FLAG (see Ozkan et al 2013). (**A**) Schematic of Drosophila brain with the subesophageal ganglion region boxed. This was the only brain region where consistent binding, with staining observed. This region is shown in the confocal images in B and D. (**B**)The genotype is *UAS-nuclear mCherry; fru P1-Gal4 (fru Pl>nuclearmcherry)*. Confocal images (40X projections) of the subesophageal ganglion region of 48 hour pupae and 0-24 hour adult male and females are shown. Binding of the Drp/DIP (green) was performed on live, dissected tissue incubated with the S2 culture media containing Dpr/DIP extracellular regions. The tissue was then washed, fixed and stained with anti-FLAG antibodies, followed by anti-mouse-Alexa 488 (green). The nuclear mCherry signal (red) is a marker for *fru P1*-expressing neurons. (**C**) The right part of each panel shows the number of cells with green Drp/DIP signal (Y axis is number of cells). The left part of each panel shows the number of cells with both green Drp/DIP and red *fru P1>nuclear mCherry* signal divided by the total number of green cells (Y axis label is the number of cells with red and green/number of green cells. Significant differences between males and females are indicated by (*). The numbers of cells that are co-expressing both proteins show significant sexual dimorphism at both time points with more co-localization in males compared to females. (**D**) The genotype is *fru^4-40^/fru^P14^*, which is a transheterozygous allele combination that is null for *fru P1*. Live staining was performed at 48 hour pupae and 0-24 hour adults, as in (**B**). (**E**) The left part of each panel shows the number of green cells detected in wild type (WT; from **C**). The right part of each panel shows the number of cells detected in *fru^4-40^/fru^P14^* in **D**. Astericks above the box plot indicates significant differences between WT and *fru^4-40^/fru^P14^* for each sex (blue indicates male comparisons, red indicates female comparisons). Black astericks below the box plots indicate differences between males and females for the *fru^4-40^/fru^P14^* analysis, at each stage. The astericks indicate: * p < 0.05, ** p < 0.01,*** p < 0.001 for student’s *t-test*. The box plot shows the first and third quartiles, with the whiskers showing the showing the minimum and maximum. Line in the box plot is the median. For all analyses, n>15 brains.

**Supplemental Figure 2.**
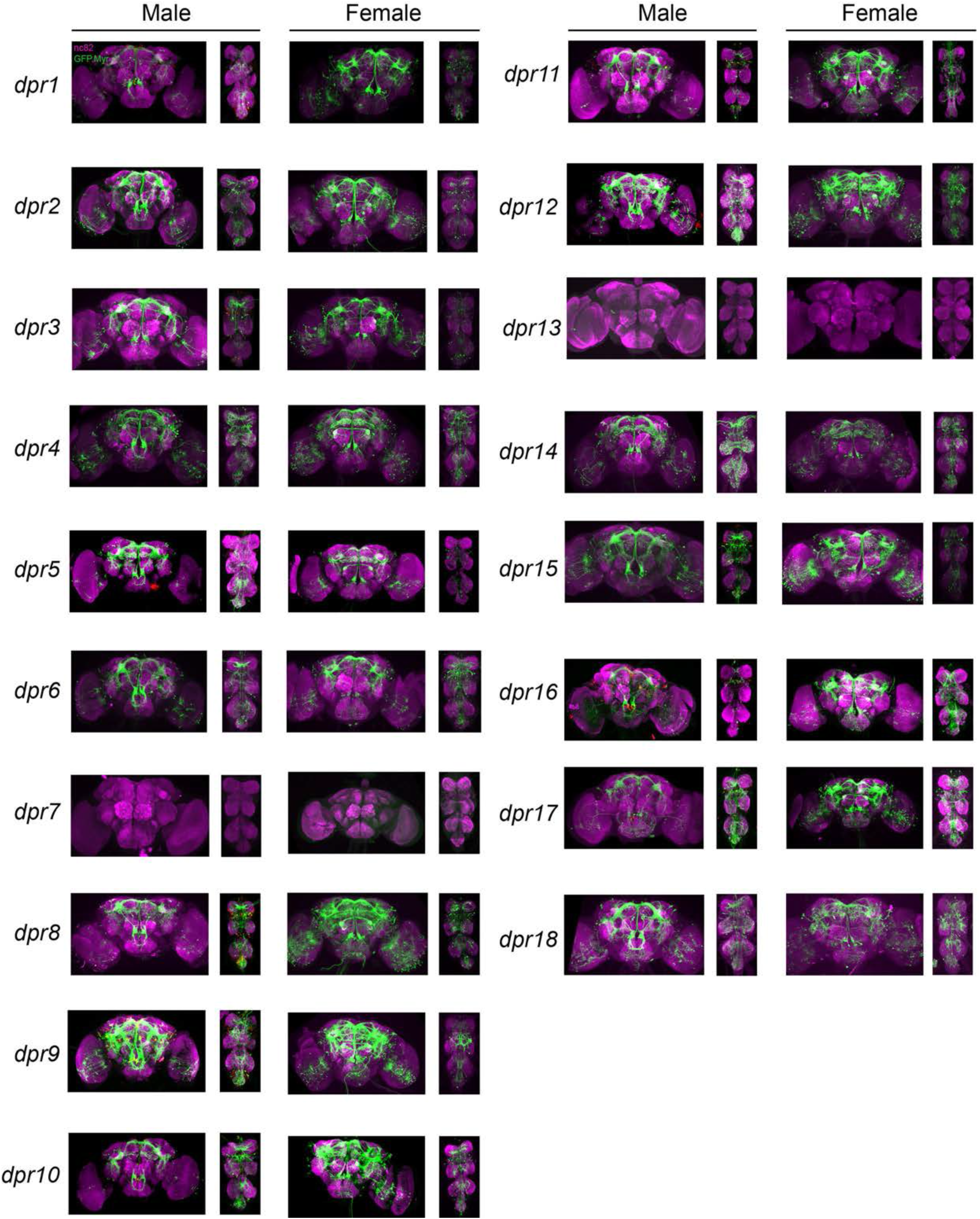
Visualization of *fru P1* and *dpr* intersecting neurons. Maximum intensity projections of brain and ventral nerve cord tissues from 0-24 hour old male and female flies. As performed in Figure 2-3.

**Supplemental Figure 3.**
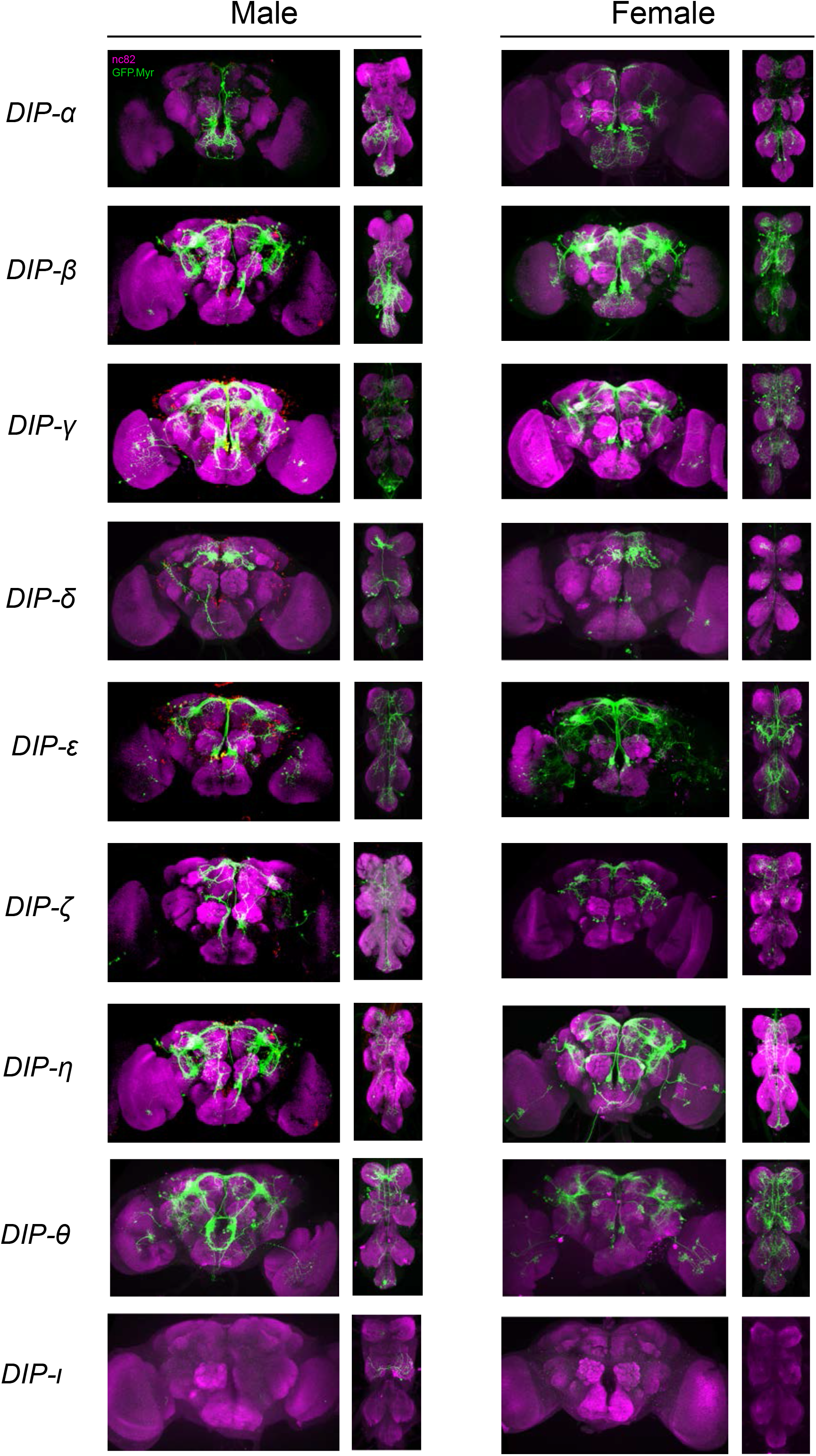
Visualization of *fru P1* and *DIP* intersecting neurons. Maximum intensity projections of brain and ventral nerve cord tissues from 0-24 hour old male and female flies. As performed in Figure 2-3.

**Supplemental Figure 4.**
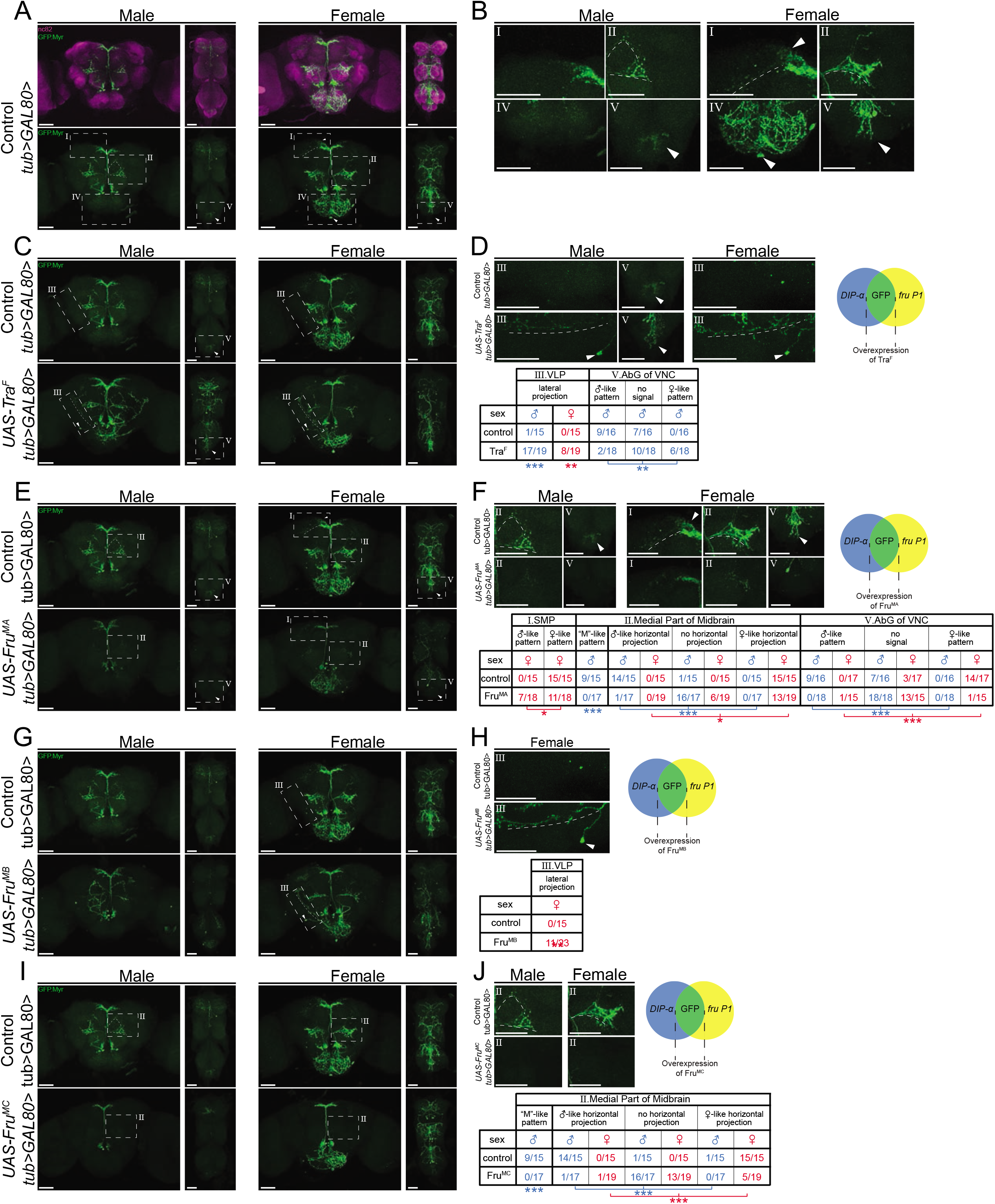
Sex hierarchy perturbation in only *fru P1 ∩ DIP-α* neurons by adding *tub>GAL80>* transgene. FLP expression, driven by *fru P1*, is required to remove GAL80 transgene, with the consquence that Gal4 is only active in intersecting neurons. The addition of *tub>GAL80>* transgene results in lower GFP amounts in both experimental and control, perhaps due to perdurance of GAL80.

**Supplemental Figure 5.**
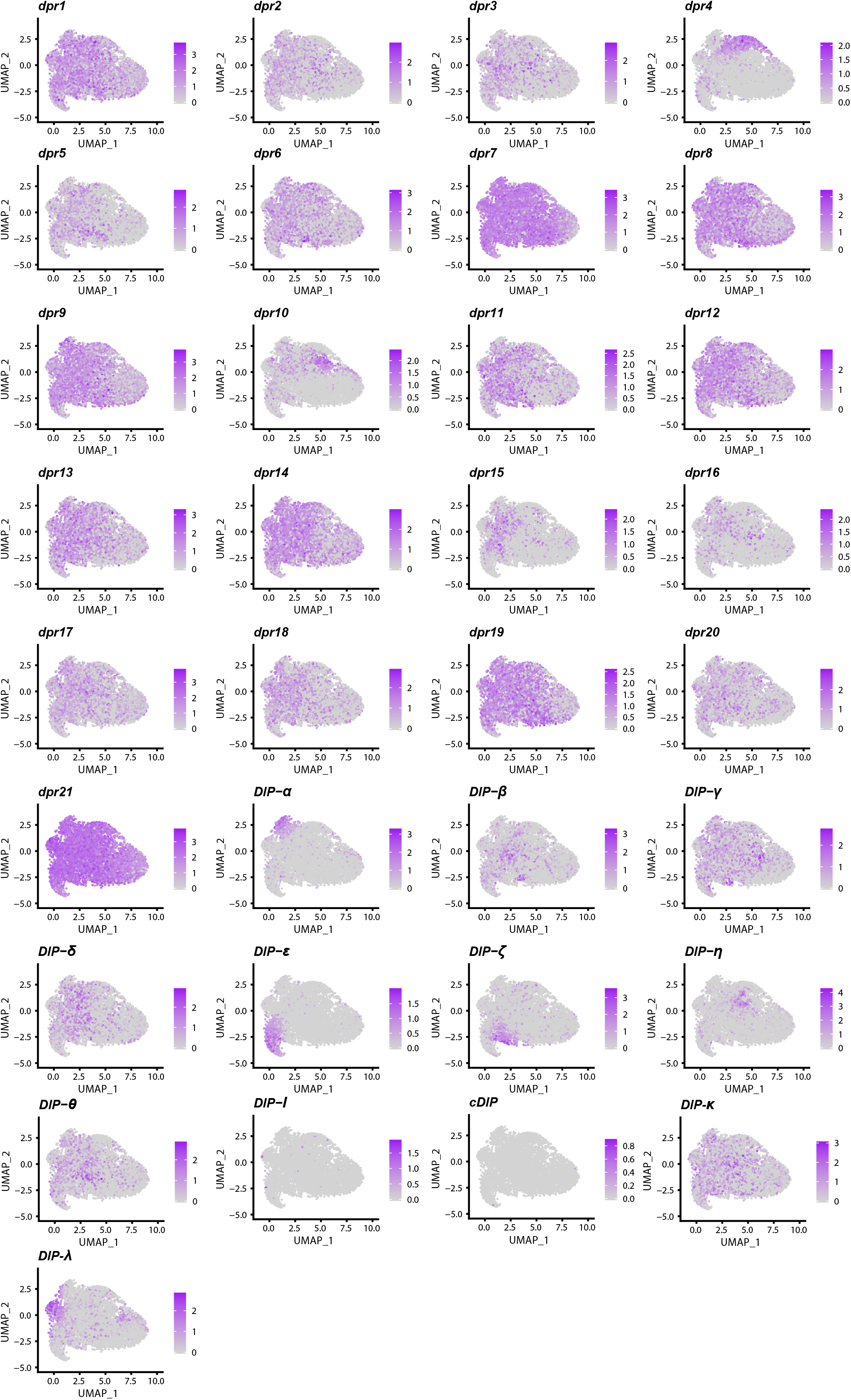
Expression of all *dpr/DIPs* in 48hr APF male *fru P1*-expressing single cells. *dpr/DIP* expression visualization in the UMAP-clustered cell (see **Figure 6A**). *dpr* or *DIP*-espressing cells are labeled purple and color intensity is proportional to log normalized expression level shown in legend nex to each UMAP panel.

